# Reprogramming Alveolar Macrophage Responses to TGF-β Reveals CCR2^+^ Monocyte Activity that Promotes Bronchiolitis Obliterans Syndrome

**DOI:** 10.1101/2022.01.27.478090

**Authors:** Zhiyi Liu, Fuyi Liao, Jihong Zhu, Dequan Zhou, Gyu Seong Heo, Hannah P. Leuhmann, Davide Scozzi, Antanisha Parks, Ramsey Hachem, Derek Byers, Laneshia K. Tague, Hrishikesh S. Kulkarni, Marlene Cano, Brian W. Wong, Wenjun Li, Howard J Haung, Alexander S. Krupnick, Daniel Kreisel, Yongjian Liu, Andrew E. Gelman

## Abstract

Bronchiolitis obliterans syndrome (BOS) is a major impediment to lung transplant survival and is generally resistant to medical therapy. Extracorporeal photophoresis (ECP) is an immunomodulatory therapy that shows promise in stabilizing BOS patients but its mechanisms of action are unclear. In a mouse lung transplant model, we show that ECP blunts alloimmune responses and inhibits BOS through lowering airway TGF-β bioavailability without altering its expression. Surprisingly, ECP-treated leukocytes are engulfed primarily by alveolar macrophages (AM), which become reprogrammed to become less responsive to TGF-β and reduce TGF-β bioavailability through secretion of the TGF-β antagonist Decorin. In untreated recipients, high airway TGF-β activity stimulates AM to express CCL2 leading to CCR2^+^ monocyte-driven BOS development. Moreover, we find TGF-β receptor 2-dependent differentiation of CCR2^+^ monocytes is required for the generation of monocyte-derived AM, which in turn promote BOS by expanding tissue-resident memory CD8^+^ T cells that inflict airway injury through Blimp-1-mediated Granzyme B expression. Thus, through studying the effects of ECP, we have identified an AM functional plasticity that controls a TGF-β-dependent network, which couples CCR2^+^ monocyte recruitment and differentiation to alloimmunity and BOS. Alveolar macrophage plasticity can be harnessed to prevent Bronchiolitis Obliterans Syndrome.

## INTRODUCTION

BOS is the most common form of chronic lung allograft dysfunction (CLAD) and the leading cause of rejection after the first year of transplantation (*1*). The major pathological hallmark of BOS is the appearance of obliterative bronchiolitis (OB), characterized by peribronchiolar and transluminal fibrotic lesions that restrict airflow (*2*). OB can also be observed in non-lung transplant settings, such as patients suffering from graft-versus-host disease or autoimmune diseases (*3*). The risk of BOS development is linked to non-alloimmune stressors that can cause bronchial injury. Club cells play a key role in bronchiolar repair through their capacity to self-renew and differentiate into goblet and ciliated cells (*4*). Previous work has shown that BOS patients have club cell dysfunction or loss (*5*). We have recently developed a lung transplant model of BOS that is triggered by the partial depletion of club cells (*6*). This model utilizes FVB (H-2^q^) donor lungs encoding three transgenes (3T-FVB); a reverse tetracycline activator gene driven by the club cell secretory protein promoter, a Cre recombinase gene under the control of the reverse tetracycline activator and a lox-P activated diphtheria toxin A gene. When 3T-FVB lungs are transplanted into immunosuppressed major histocompatibility complex (MHC)- mismatched C57BL/6 (B6; H-2^b^) recipients, club cell depletion after transient doxycycline (DOX) ingestion results in bronchiolar injury and the development of severe OB lesions. Importantly, lymphocyte-mediated immune responses against allo- and autoantigens, known target antigens in BOS subjects, develop in this model (*5–7*). However, in syngeneic 3T lung transplant recipients, club cell depletion-mediated bronchiolar injury is repaired and fails to produce OB lesions (*6*).

ECP is an autologous cell-based immunotherapy where apheresed peripheral blood leukocytes are treated with 8-methoxypsoralen and ultraviolet A radiation prior to re-infusion. ECP has been used for a wide variety of chronic inflammatory disorders and is currently being investigated as treatment for BOS (*8–10*). Although randomized double-blind trials have yet to be completed, there is accumulating evidence that ECP improves lung function or prevents its decline (*11*). Additionally, beneficial responses to ECP have been shown to coincide with the reduction of circulating allo- and autoantibodies (*12*). Previous studies have reported ECP increases TGF-b protein expression but whether it regulates bioavailability is unclear (*13*). Although TGF-b is required to maintain tissue homeostasis and helps promote the resolution of inflammation (*14*) it is also a potent mediator of tissue fibrosis (*15*). TGF-b is secreted complexed to a latent activating peptide and a latent TGF-b binding protein, which adheres it to the extracellular matrix (*16*). Following tissue injury, TGF-b can become active through its release from latent complex proteins by a diverse set of factors that include alpha integrin αvβ_5_, metalloproteases and cathepsins (*17*). However, even after becoming active, TGF-b can be re-regulated by soluble leucine rich proteoglycans such as Decorin, which bind to TGF-β to prevent engagement with its receptor (*18*).

AM play a critical role in maintaining distal airway homeostasis through promoting host defense and performing surfactant catabolism (*19*). In quiescent lungs, the AM compartment is nearly entirely composed of self-renewing tissue-resident AM (TR-AM) that develop during embryogenesis (*20*). However, during pulmonary inflammation TR-AM levels fall coincident with the generation of monocyte-derived AM (Mo-AM) (*21*). In comparison to TR-AM, the specific requirements for Mo-AM development are less defined. Several reports show that Mo-AM are derived from bone marrow-derived cells, but have not directly addressed whether this AM subset arises from CCR2^+^ monocytes (*22, 23*). CCR2 expression on monocytes is critical for trafficking into inflamed lungs in response to the chemokine CCL2 (*24, 25*). CCR2 expression and Mo-AM have been shown to drive bleomycin-induced pulmonary fibrosis raising the possibility that CCR2^+^ monocytes drive pulmonary fibrotic disease through differentiation into Mo-AM (*21, 26*). The prevailing view is that AM are poor antigen presenting cells that function primarily to enforce airway tolerance (*27*). However, while controversial, some recent observations have indicated that AM are capable of promoting the effector activity of tissue-resident memory CD8^+^ T cells (T_RM_) (*28, 29*). T_RM_ differ from other memory subsets because they do not recirculate and develop in peripheral tissues under the instruction of locally derived cues, such as TGF-β (*30, 31*). Nevertheless, they share some properties with effector memory cells such as the expression of the transcription factor *Pdrm1* (Blimp-1), which drives the expression of Granzyme b (Gzmb) (*32*).

Here, by studying the effects of ECP we uncovered immune mechanisms that promote BOS after lung transplantation. ECP inhibits BOS through reducing AM responses to TGF-β and lowering intragraft TGF-β bioavailability by inducing Decorin expression. In untreated recipients, high intragraft TGF-β activity stimulates AM to express CCL2 that in turn drives CCR2^+^ monocyte allograft recruitment and promotes TGF-β receptor 2-dependent CCR2^+^ monocyte differentiation into Mo-AM. We observe stable interactions between T_RM_ and AM by intravital two-photon microscopy and show that Mo-AM reactivate T_RM_ through donor-antigen presentation. Finally, we demonstrate that Mo-AM promote BOS through stimulating the expansion of Blimp-1^+^Gzmb^hi^ T_RM_.

## RESULTS

### ECP inhibits BOS and blunts lymphocyte recognition of transplant antigens

To analyze the immunoregulatory effects of ECP we utilized mouse donor lungs (3T-FVB) that when transplanted into immunosuppressed B6 recipients develop BOS following doxycycline (DOX)-mediated ingestion to induce diptheria toxin expression in club cells (*6*). Following DOX ingestion, lung recipients received intravenous infusions of ECP-treated B6 leukocytes at three-day intervals and allografts were analyzed for histological appearance and lymphocyte activation on post-operative day (POD) 16 (**Fig. 1A**). In contrast to saline vehicle-treated mice, recipients that received ECP had significantly less peribronchiolar inflammation, largely devoid of OB lesions and were significantly able to regenerate club cells (**Fig. 1B-F**). ECP treatment also reduced and intragraft hydroxyproline content and neutrophilia (**Fig. 1G, H**). Analysis of allograft infiltrate showed a lower percent abundance of IL-17A^+^ CD4 ^+^ and IFN-γ^+^ CD8^+^ T cells (**Fig. 1I, Fig. S1**). We next assessed the effects of ECP on transplant-antigen recognition by lymphocytes (**Fig. 1J-M**). When compared to saline-treated recipients, CD4^+^ and CD8^+^ T cells from ECP-treated recipients had reduced CD4^+^ T cell-mediated IL-17 production and less CD8^+^ T cell-mediated IFN-γ production following re-stimulation with donor antigens. Subjects that develop BOS have been reported to have lymphocytes that recognize the lung self-antigens Collagen V (Col V) and k-alpha tubulin (Kα1T) (*7, 12*). ECP-treated allograft resident CD4^+^ T cells when challenged with Col V or Kα1T peptides expressed less IL-17A when compared to saline-treated recipients. Finally, and in line with previous clinical observations (*12*), ECP reduced the serum levels of donor-specific antibodies (DSA). These results demonstrate that ECP inhibits transplant antigen-specific responses and reduces BOS severity.

**Figure 1:**
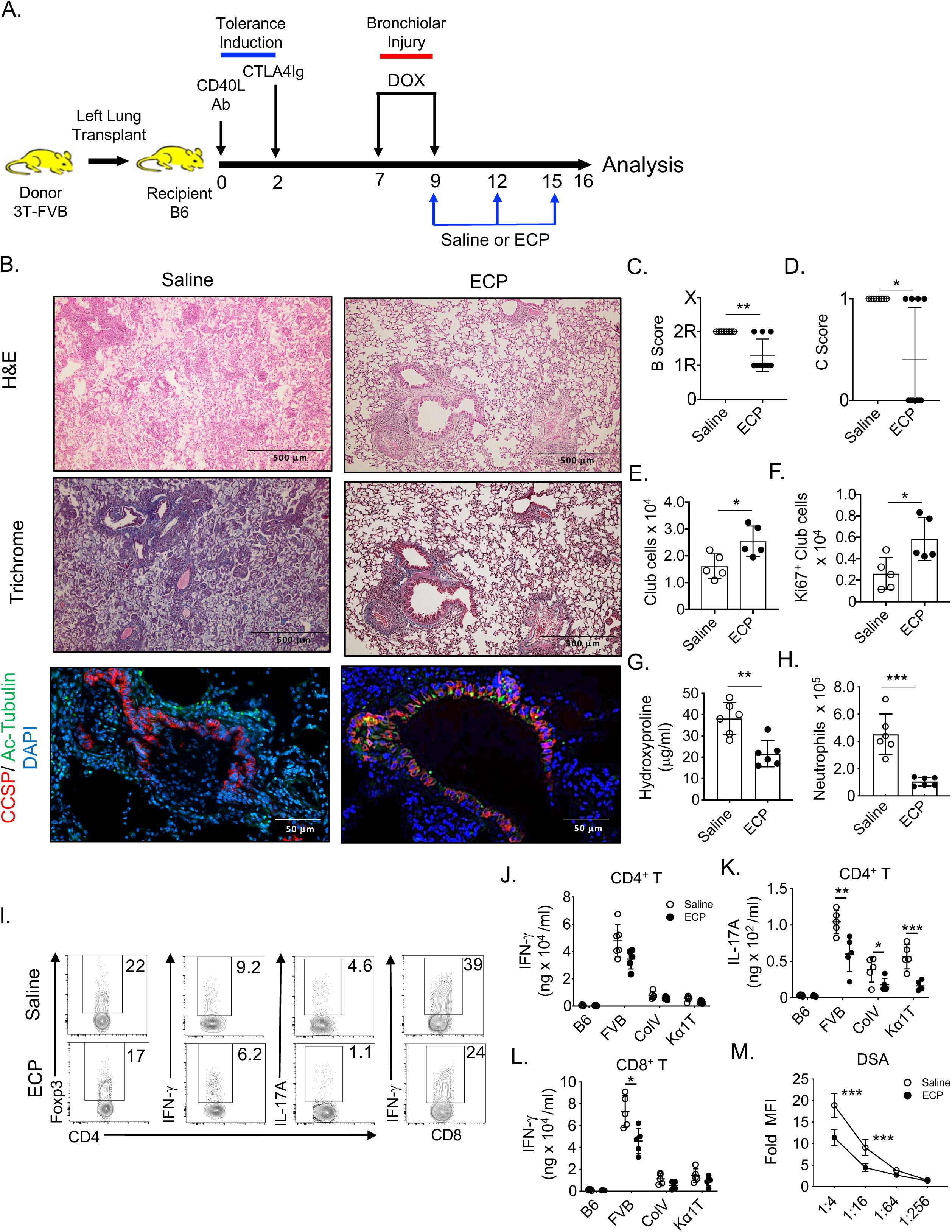
ECP prevents BOS and lymphocyte recognition of transplant antigens. **(A)** 3T-FVB left lungs were transplanted into C57Bl/6 (B6) and treated with CD40L Abs (POD 0) and CTLA4Ig (POD 2) to establish allograft tolerance. Between POD 7 and 9 recipients ingest doxycycline (DOX), receive saline or ECP-treated B6 leukocytes on POD 9, 12 and 15 and euthanized on POD 16. (**B**) Representative allograft H&E, Trichrome and Club cell secretory protein (CCSP)/Ac-tubulin Ab staining (N ≥ 9/group). (**C, D)** Allografts scored for airway inflammation (B score) where 0 = none, 1R= low grade, 2R = high grade and X = ungradable and the (C score) presence (1) or absence (0) of OB lesions. Intragraft (**E**) total and (**F**) Ki67^+^ club cell numbers, (**G**) Hydroxproline content, (**H**) neutrophilia. (**I)** Representative FACS plots of the percent abundance for indicated T lymphocyte lineages. (**J-L**) T cell antigen specificity measured by IFN-*γ*and IL-17A production following stimulation with splenocytes isolated from B6 (syngeneic antigens), FVB (donor antigens) or B6 mice pulsed with lung self-antigens Col V and K*α*1T. **(M)** DSA (IgM) serum reactivity against FVB CD19^+^ cells at indicated dilutions. Bars represent means ± S.D where *p < 0.05; **p < 0.01, *** p < 0.001.

### ECP reprograms AM to inhibit intragraft TGF-β bioavailability

Given reports that ECP stimulates TGF-β production (*13*), we measured protein levels of all three TGF-β isoforms in the bronchioalveolar lavage fluid (BALF) and the peripheral serum of saline- and ECP-treated 3T-FVB as well as 2T-FVB lung recipients, which maintain established tolerance despite DOX ingestion as they do not undergo bronchiolar injury due to the lack of the *lox-P* activated diphtheria toxin A gene (*6*) (**Fig. 2A**). Relative to 2T-FVB lung recipients, BALF TGF-β1 levels were significantly elevated in saline- and ECP-treated 3T-FVB recipients. However, BALF TGF-β1 accumulation was not significantly different between saline- and ECP- treated 3T-FVB recipients. Additionally, BALF and circulating TGF-β2 and TGF-β3 levels were either undetectable or were expressed in very modest quantities in all lung recipients. Because TGF-β bioavailability is highly regulated (*17*), we next assessed the ability of lung recipient BALF and peripheral serum to induce TGF-β-receptor signaling using a SMAD2/3 reporter cell line (**Fig. 2B**). BALF from ECP-treated recipients had significantly less TGF-β activity when compared to saline-treated 3T-FVB hosts. In contrast, no significant differences in TGF-β activity were detected in the peripheral serum from saline-, ECP-treated 3T-FVB or 2T-FVB lung recipients. In light of recent observations that immunoregulatory circuits act locally within lung transplants to control tolerance (*33*), we next analyzed ECP-treated leukocyte trafficking to 3T-FVB allografts undergoing BOS pathogenesis. ECP-treated cells were labeled with a fluorescent dye and assessed for engulfment by 3T-FVB allograft CD11b^+^ phagocytes two hours after intravenous injection (**Fig. 2C, Fig. S2A**). Approximately three quarters of ECP-treated leukocytes were engulfed by Mo-AM and TR-AM, which were differentiated by expression of CD45.1 and CD45.2 in the donor and recipient, respectively.

**Figure 2:**
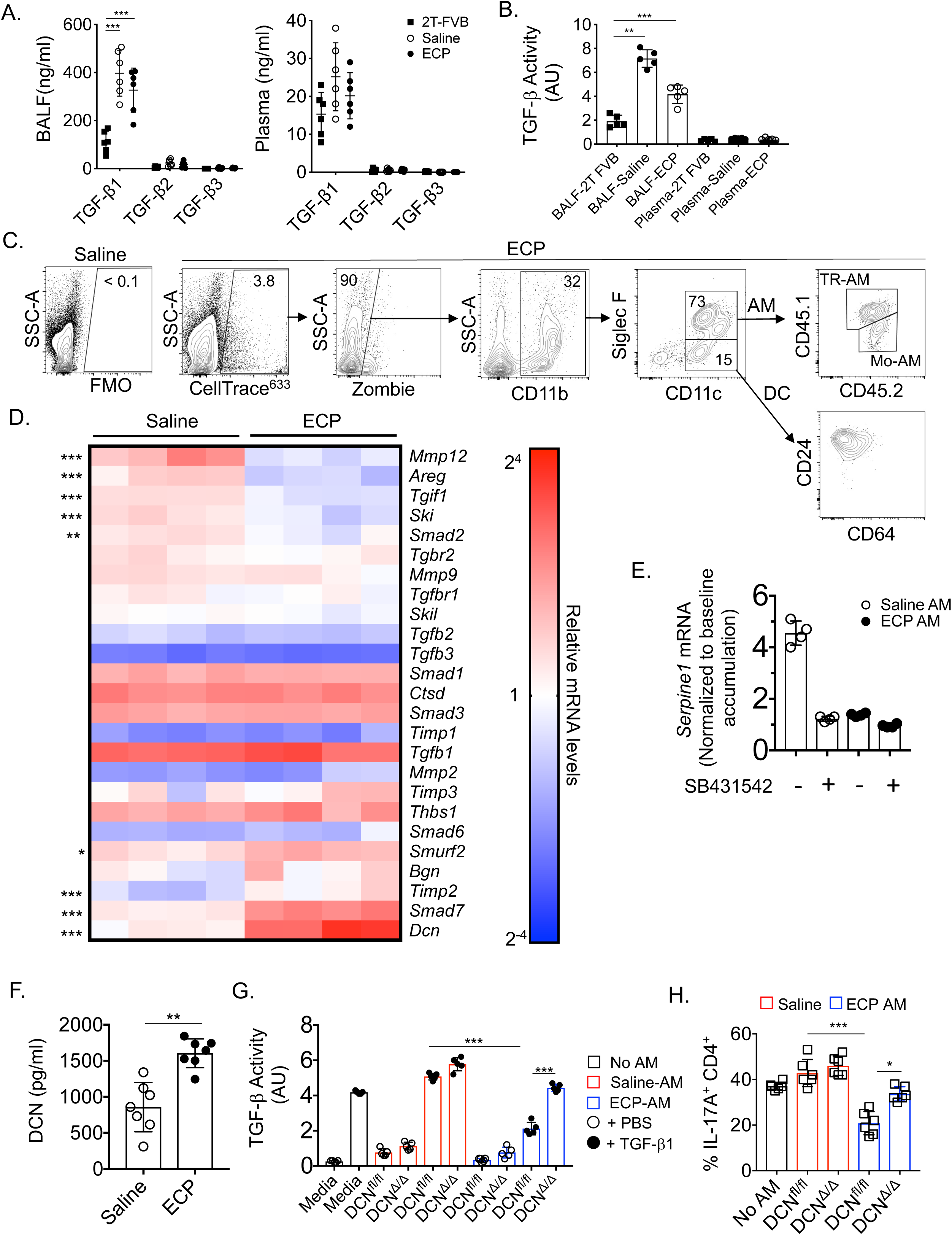
ECP reprograms AM to antagonize TGF-β bioavailability. POD 16 2T and 3T FVB allograft **(A)** BALF and plasma analyzed for TGF-β isoform protein content by ELISA or **(B)** activity with a HEK 293 SMAD 2/3 luciferase reporter cell line. AU represent arbitrary luciferase units. **(C)** CellTrace^633^ labeled ECP-treated leukocytes injected into 3T FVB allograft and analyzed for uptake by intragraft CD11b^+^ phagocytes. Data shown are one set of representative FACS plots from four individually conducted experiments. (**D**) Heat map of saline- and ECP-treated POD 16 3T-FVB allografts AM transcript levels of TGF-β signaling and fibrosis related gene targets normalized to the macrophage housekeeping gene *Sxt5a*. **(E)** Fold accumulation of TGF-β-induced AM *Serpine1* mRNA accumulation in the presence or absence of 10 μM SB43152 or vehicle (DMSO). Data shown is normalized to baseline levels (non-TGF- β-treated DMSO-pretreated controls). **(F)** Saline and ECP-treated AM were cultured overnight and analyzed by ELISA for DCN secretion (N = 8/group). **(G)** TGF-β activity measurements of enriched supernatants from saline- or ECP treated DCN^Δ^*^/^*^Δ^ and DCN^fl/fl^ AM cultured with or without 10 ng/ml TGF-β1. **(H)** Plate bound CD3ε and CD28 Ab-mediated stimulated B6 naïve CD4^+^ T cell were cultured for 4 days assessed for differentiation into IL-17A^+^ CD4^+^ T cells in the presence or absence of indicated AM conditioned supernatants added in a 1:1 ratio to T_h_17 polarization medium that contains 10 ng/ml TGF-β1. Bars represent means ± S.D where *p < 0.05; **p < 0.01, *** p < 0.001.

Given that AM uptake of ECP-treated leukocytes was linked to lower airway TGF-β bioavailability we next analyzed 3T-FVB allograft AM transcript levels of 25 genes reported to control TGF-β responses and activation in lung macrophages (*23, 34, 35*) (**Fig. 2D**). Nine transcripts were found to be differentially regulated by ECP. For example, several genes that inhibit TGF-β signaling, *Dcn*, *Smad*7 and *Smurf2,* were significantly upregulated in ECP-treated AM (*36*). Conversely, factors that promote the activation of latent TGF-β such as *Mmp13* and *Areg* (*37*) were downregulated by ECP. Interestingly, genes that regulate TGF-β signaling such as *Tgif1* and *Ski* were also downregulated in ECP-treated AM suggesting a compensatory response due to a lack of homeostatic TGF-β receptor signaling (*23*). To further confirm these observations, we analyzed TGF-β1-mediated responses of two known TGF-β expression targets, *Serpine1* (*38*) and the TGF-β-activating integrin αvβ5 (*39*), in saline- and ECP-treated AM (**Fig. 2E, Fig. S2B**). Unlike in ECP-treated AM, TGF-β1 induced *Serpine1* mRNA accumulation and αvβ5 protein upregulation in saline-treated AM. To determine if these reductions were TGF-β signaling dependent, ECP- and saline-treated AM were also pre-incubated with the TGF-β receptor inhibitor SB431542 (*40*) prior to stimulation with TGF-β1. SB431542 addition to saline-treated AM inhibited TGF-β1-mediated *Serpine1* and αvβ5 gene induction to levels nearly comparable to TGF-β1-stimulated ECP-treated AM.

Since *Dcn* (DCN) was the most differentially regulated transcript in our analysis we measured its secretion (**Fig. 2F**). DCN secretion was significantly higher in ECP-treated AM when compared to saline-treated AM. We further sought to determine if ECP-treated AM regulates TGF-β activity in DCN-dependent manner. For this purpose, we generated *Lyz^Cre/+^ Dcn ^fl/fl^* (DCN^Δ/Δ^) mice and tested the ability of conditioned supernatants from ECP-treated DCN^Δ/Δ^ and wildtype control *Dcn ^fl/fl^* (DCN^fl/fl^) AM to stimulate TGF-β signaling activity (**Fig. 2G**). Supernatants from ECP-treated DCN^fl/fl^ AM sharply reduced TGF-β activity when compared to ECP-treated DCN^Δ/Δ^ AM or saline-treated DCN^fl/fl^ AM. Notably, alterations in TGF-β activity were most apparent when TGF-β1 was added to cultures indicating ECP-treated AM primarily target TGF-β activity generated by exogeneous sources. TGF-β drives T_h_17 generation from naive CD4^+^ T cells (*41*) and also promotes T_h_17 lineage stability (*42*). Given that ECP treatment reduces intragraft IL-17A^+^ CD4^+^ T cell accumulation, we analyzed the effects of saline- and ECP-treated AM conditioned supernatants on T_h_17 cell development (**Fig. 2H, Fig. S3**). Differentiation of naïve CD4^+^ T cells into IL-17A^+^ CD4^+^ T cells was significantly impeded by ECP-treated DCN^fl/fl^ AM when compared to ECP-treated DCN^Δ/Δ^ AM or saline-treated AM irrespective of DCN expression. Collectively, these data show that ECP induces AM to become unresponsive to TGF-β signals and to reduce local TGF-β bioavailability.

### ECP-mediated inhibition of BOS is dependent on AM Decorin expression

To determine if AM DCN expression is required for ECP-mediated attenuation of BOS we first reconstituted the donor allograft AM compartment with DCN^Δ/Δ^ and DCN^fl/fl^ AM. To this end, we administered clodronate liposomes intratracheally into 3T-FVB mice just prior to transplantation to deplete donor wildtype TR-AM (*21*). Importantly, clodronate treatment led to an approximately 95% reconstitution of the AM compartment with recipient-derived AM but did not prevent the induction of immunosuppression-mediated acceptance or spontaneously induce BOS lesions (**Fig. S4A-C**). However, following bronchiolar injury, ECP was ineffective at attenuating BOS and failed to reduce IL-17A^+^ CD4^+^ or IFN-γ^+^ CD8^+^ T cell intragraft accumulation in DCN^Δ/Δ^ recipients (**Fig. 3A-D**). In contrast, ECP-treated wildtype DCN^fl/fl^ recipients were protected from BOS and had lower BALF TGF-β activity when compared to ECP-treated DCN^Δ/Δ^ recipients. Because DCN is reported to interact with other growth factors that regulate inflammation (*43*), it remained possible that our observed effects on TGF-β activity were unrelated to inhibiting BOS development. For this purpose, we tested the effects of TGF-β antibody blockade on BOS development (**Fig. 3E-G**). TGF-β neutralizing antibodies were administered intratracheally into 3T-FVB lungs that were transplanted into B6 recipients and treated with DOX. T cell activation and BOS development were inhibited, comparable to our observations after ECP treatment. Overall, these data indicate that AM-mediated regulation of TGF-β bioavailability controls BOS pathogenesis.

**Figure 3:**
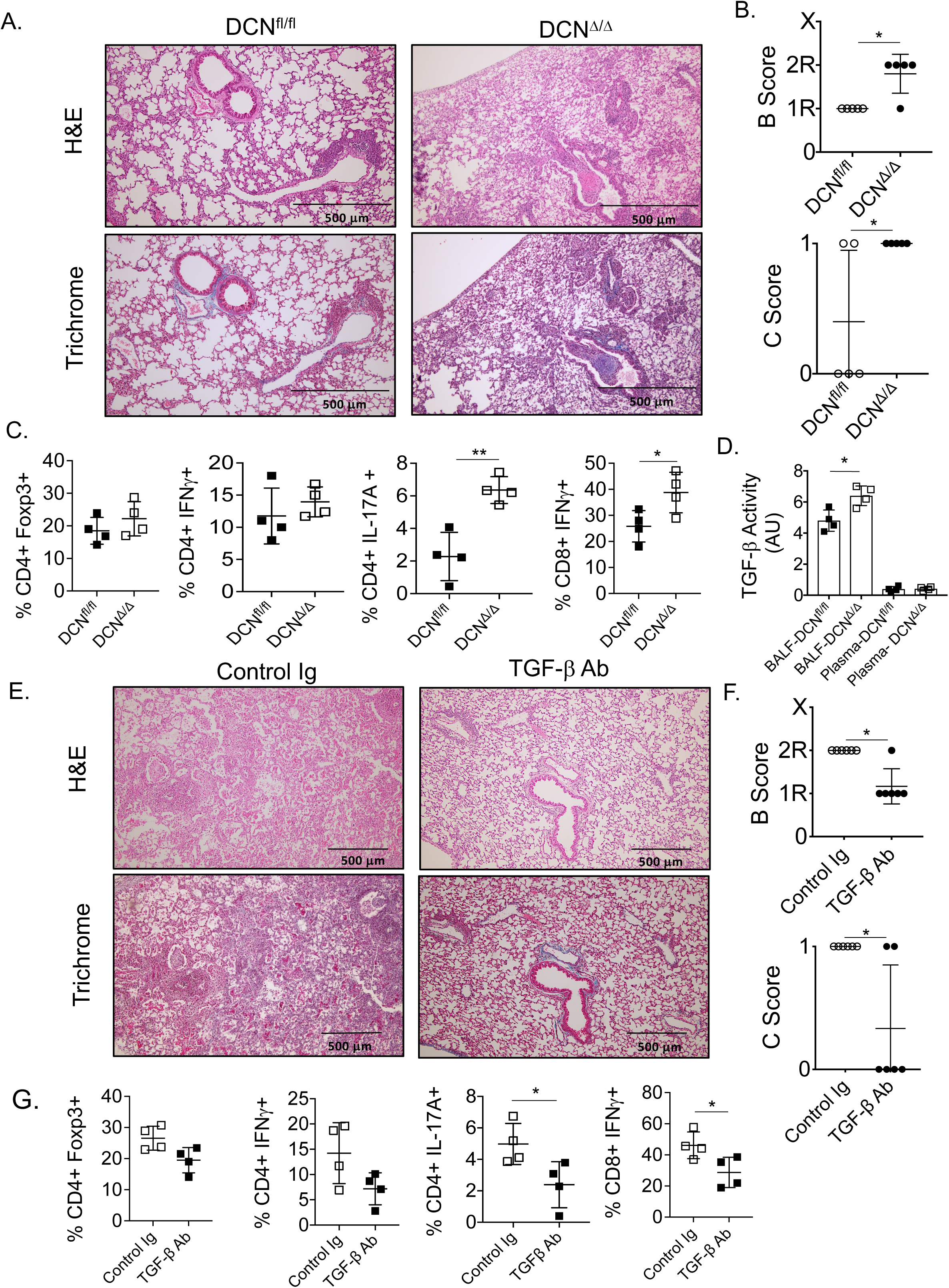
ECP-mediated inhibition of BOS targets TGF-β activity. Prior to transplantation 3T-FVB allografts are treated on POD -1 with intratracheal clodronate liposomes (100 μl) to deplete airway AM and then transplanted into ECP-treated DCN^Δ^*^/^*^Δ^ and DCN^fl/fl^ recipients and then evaluated for intragraft inflammation on POD 16 as shown by (**A**) a representative image of H&E and Trichrome staining (N=4/group), (**B**) airway inflammation and lesion grading, (**C**) intragraft T cell activation and (**D**) BALF and circulating plasma TGF-β activity. B6 recipients of 3T FVB allografts received intratracheal Mouse IgG or TGF-β Ab (75 μg/100 ul PBS) on POD 7 and on POD 16 analyzed for intragraft inflammation as shown by (**E**) a representative image of H&E and Trichrome staining (N=4/group), (**F**) airway inflammation and lesion grading and (**G**) intragraft T cell activation. Bars represent means ± S.D where *p < 0.05; **p < 0.01.

### Infiltrating CCR2^+^ monocytes promote BOS

Given previous reports that recipient CCR2 deficiency prevents fibrosis in mouse non-vascularized tracheal allografts (*44*), we next set out to assess ECP-mediated changes in CCR2 expression within lung allografts. To this end, we imaged ECP-treated lung recipients using a positron emission tomography (PET) purposed radiotracer, ^64^Cu-DOTA-ECL1i, which specifically recognizes the extracellular loop number one of CCR2 and is under current clinical evaluation for non-invasive diagnosis of idiopathic pulmonary fibrosis (*45*). When compared to untreated 3T-FVB allografts that develop BOS, we observed a sharp decrease in CCR2 activity in the allografts when recipients were treated with ECP (**Fig. 4A, B**). We next determined if CCR2^+^ monocytes are required for BOS development. For this purpose, we transplanted 3T-FVB lungs into CCR2^+^ monocyte depleter recipients, which encode the diptheria toxin receptor under the control of the CCR2 promoter (CCR2^DTR^) (*46*). We also inhibited the activity of CCL2 by injecting CCL2 neutralizing antibodies into B6 recipients of 3T-FVB allografts (**Fig. 4 C-E, Fig. S5A, B**). CCR2^+^ monocyte depletion or CCL2 antibody blockade reduced the development of severe OB lesions. Additionally, we observed a significant reduction in intragraft IFN-γ^+^ CD8^+^ T cells when compared to control conditions. Collectively, our data show that CCR2^+^ monocyte allograft infiltration promotes BOS.

**Figure 4:**
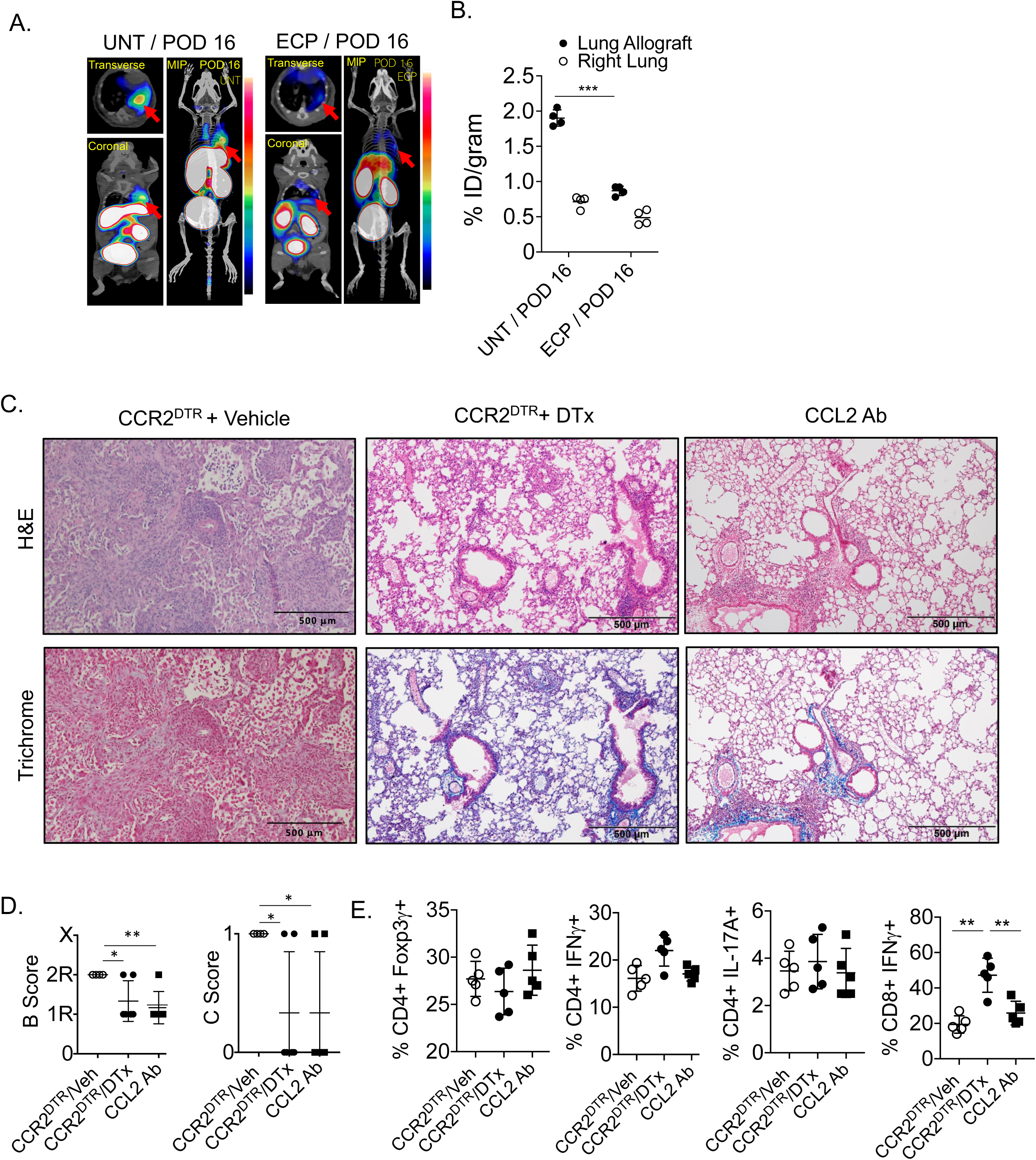
CCR2^+^ monocytes drive BOS. (A) Representative dynamic ^64^CuDOTA-ECL1i PET/CT images scans of untreated and ECP-treated 3T-FVB allografts (red arrows) with (B) right native lung and allograft probe uptake quantitation shown as percent injected dose per gram (%/ID/gram) of tissue (N=4/group). (C) 3T FVB allografts of CCR2^DTR^ recipients received 10 ng/g i.v. of diptheria toxin on POD 6 and 11 and B6 recipients of 3T FVB allografts received 200 ug i.v. of CCL2 neutralizing Ab on POD 6, 9 and 12. Both recipients were euthanized on POD 16 and assessed for intragraft inflammation by (**C**) representative H&E and Trichrome staining (N=5/group), (**D**) airway inflammation and lesion grading and (**E**) intragraft T cell activation. Bars represent means ± S.D where *p < 0.01, *** p < 0.001.

### TGF-β stimulates AM CCL2 expression to promote Mo-AM allograft accumulation

We have recently reported that AM produce CCL2 after lung transplantation (*25*). TGF-β targets activation of AP-1 and EGR1, transcription factors that can promote CCL2 gene transcription (*47, 48*). We stimulated saline- and ECP-treated AM with TGF-β1 and measured CCL2 production (**Fig. 5A**). TGF-β1 significantly increased CCL2 expression in saline-, but not ECP-treated AM. Low molecular weight hyaluronic acid (HA), a damage-associated molecular pattern that we have shown accumulates in lung transplants with BOS (*49*), has also been demonstrated to promote CCL2 expression in a mouse AM cell line (*50*). HA addition to TGF-β1-stimulated cultures induced a synergistic increase in CCL2 expression in saline-treated AM relative to HA stimulation alone. In ECP-treated AM we observed that CCL2 expression mediated by TGF-β-HA costimulation was comparable to HA stimulation indicating a lack of a synergistic response in these cells. We next asked if AM-mediated CCL2 production during BOS pathogenesis induces CCR2^+^ monocyte allograft infiltration (**Fig. 5B**). 3T-FVB allografts were depleted of AM or treated with anti-TGF-β antibodies prior to bronchiolar injury and assessed for airway CCL2 production and numbers of recruited CCR2^+^ monocytes. Although we could detect some CCL2 production in allografts prior to bronchiolar injury, levels rose approximately sixfold following bronchiolar injury. However, CCL2 levels were substantially reduced by either AM depletion or TGF-β antibody blockade. Similarly, CCR2^+^ monocyte allograft accumulation was significantly attenuated by either treatment.

**Figure 5:**
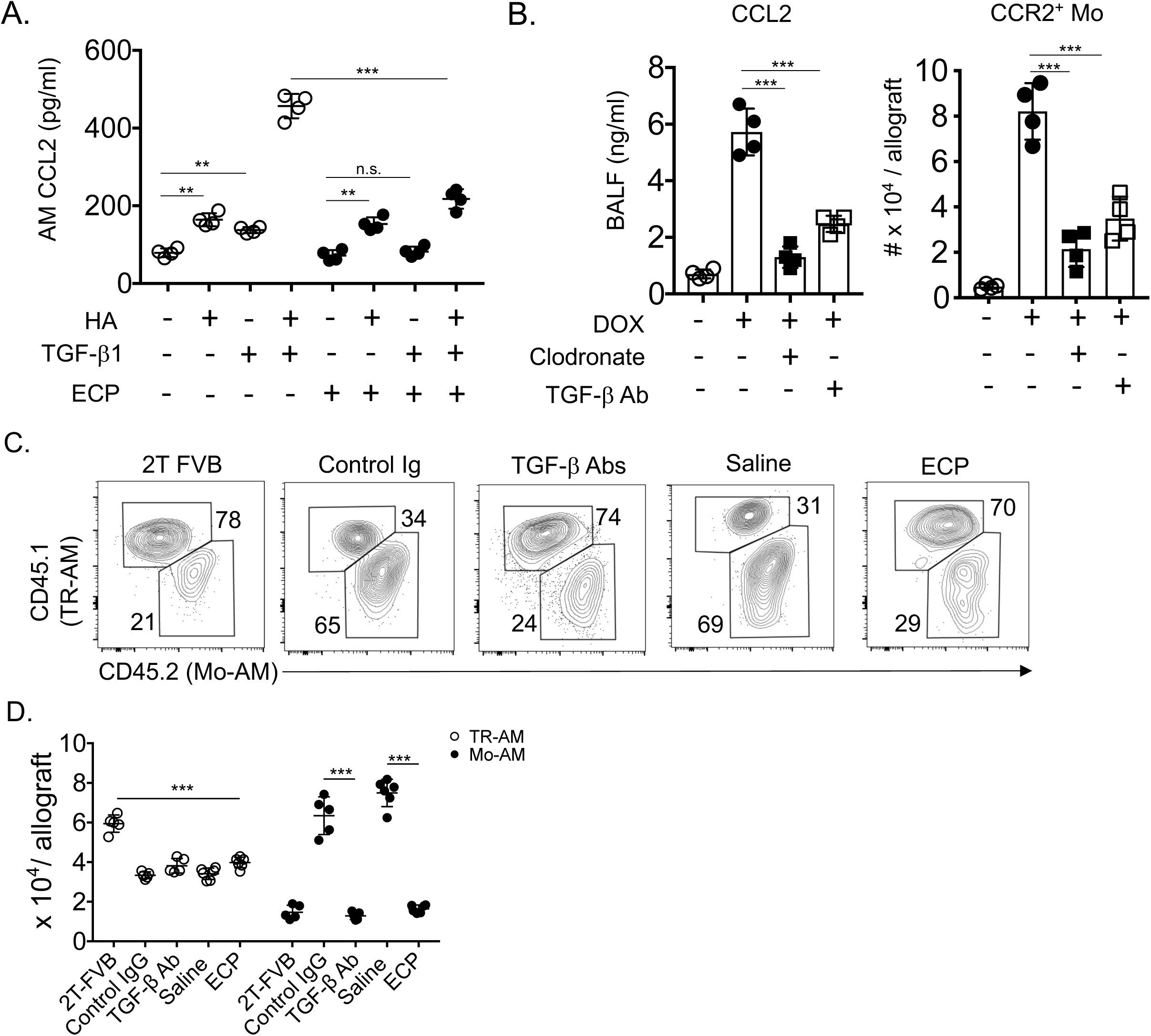
A TGF-β-AM-CCL2 expression circuit promotes Mo-AM allograft accumulation. (**A**) Untreated and ECP treated AM were stimulated with 10 ng/ml TGF-β1 and/or 1 μg/ml HA for 18 hrs and assessed for CCL2 expression by ELISA. (**B**) POD 6 3T-FVB allograft recipients were treated with intratracheal TGF-β Ab (75 μg/100 ul PBS) or clodronate liposomes (100 μl), induced to undergo bronchiolar injury (DOX) on POD 7 and assessed for BALF CCL2 expression and CCR2^+^ monocyte recruitment on POD 8. 2T-FVB and 3T-FVB allograft recipients underwent indicated treatments and were quantitated for Mo-AM and TR-AM as shown by (**C**) representative plots of relative percent abundance (N= 5/group) and (**D**) cell counts. Bars represent means ± S.D where *p < 0.05; **p < 0.01, *** p < 0.001.

Given previous observations that Mo-AM drive pulmonary fibrogenesis (*21*), we quantified Mo-AM and TR-AM in 3T-FVB allograft recipients treated with saline, ECP, control Ig or TGF-β-neutralizing antibodies (**Fig. 5C, D**). Relative to 2T-FVB allograft recipients with established tolerance, we observed a nearly uniform reduction in TR-AMs in 3T-FVB allografts irrespective of treatment. In contrast, Mo-AM levels were affected by treatments that target TGF-β bioavailability. In control Ig or saline-treated lung recipients, Mo-AM were approximately twofold more abundant than TR-AM and were approximately fourfold more abundant when compared to Mo-AM in TGF-β antibody- or ECP-treated allografts. Collectively, these data show that TGF-β induces AM to express CCL2, which in turn promotes the allograft recruitment of CCR2^+^ monocytes and Mo-AM.

### TGF-β-receptor mediated CCR2^+^ monocyte differentiation into Mo-AM leads to BOS

The correlation between CCR2^+^ monocyte and Mo-AM allograft accumulation during BOS pathogenesis raises the possibility that CCR2^+^ monocytes differentiate into Mo-AM. Interestingly, previous work has shown that TGF-β promotes AM development from bone marrow-derived cells (*23*). To determine if TGF-β drives CCR2^+^ monocyte differentiation into Mo-AM, we generated tamoxifen-inducible *Cc2r^CreERT2/+^ Tgfbr2^fl/fl^* (TGF-βR2^Δ/Δ^) mice. Tamoxifen treatment of TGF-βR2^Δ/Δ^ CCR2^+^ monocytes reduced *Tgfbr2* mRNA by nearly 80% relative to *Tgfbr2^fl/fl^* (TGF-βR2^fl/fl^) controls and inhibited TGF-β1-mediated generation of two transcripts required for AM development, *Pparg* and *Car4* (*51*) (**Fig. 6SA, B**). TGF-βR2^Δ/Δ^ recipients of 3T-FVB lungs were comparatively poor at inducing Mo-AM accumulation when compared to wildtype TGF-βR2^fl/fl^ recipients (**Fig. 6A**). Notably, the sharp reduction in Mo-AM graft accumulation in TGF-βR2^Δ/Δ^ recipients was not due to a defect in CCR2^+^ monocyte recruitment following bronchiolar injury (**Fig. 6B**). Moreover, TGF-βR2^Δ/Δ^ lung recipients were significantly protected from BOS, which was associated with a reduction in intragraft IFNγ^+^ CD8^+^ T cells (**Fig. 6C-E**). As the reduction in BOS could be explained by the inability to generate other CCR2^+^ monocyte descendants we created reporter *Ccr2^CreERT2/+^ Tgfbr2^fl/fl^ TdTomato^fl/STOP/+^* and *Ccr2^CreERT2/+^ Tgfbr2^+/+^ TdTomato^fl/STOP/+^* mice to conduct fate mapping studies. Following tamoxifen treatment, bone-marrow derived CCR2^+^ monocytes were adoptively transferred into 3T-FVB lung transplant recipients undergoing BOS pathogenesis (**Fig. 6F**). Irrespective of TGF-βR2 deletion, we detected similar numbers of CD11c^+^ descendants within allografts. However, TGF-βR2-deficient CCR2^+^ monocytes were profoundly deficient at generating Mo-AM, but iMac and CD11b^+^ DC developed independently of TGF-βR2. Collectively, these data indicate that intrinsic CCR2^+^ monocyte TGF-β signaling is required for Mo-AM development but not for the generation of other CD11c^+^ derived lineages.

**Figure 6:**
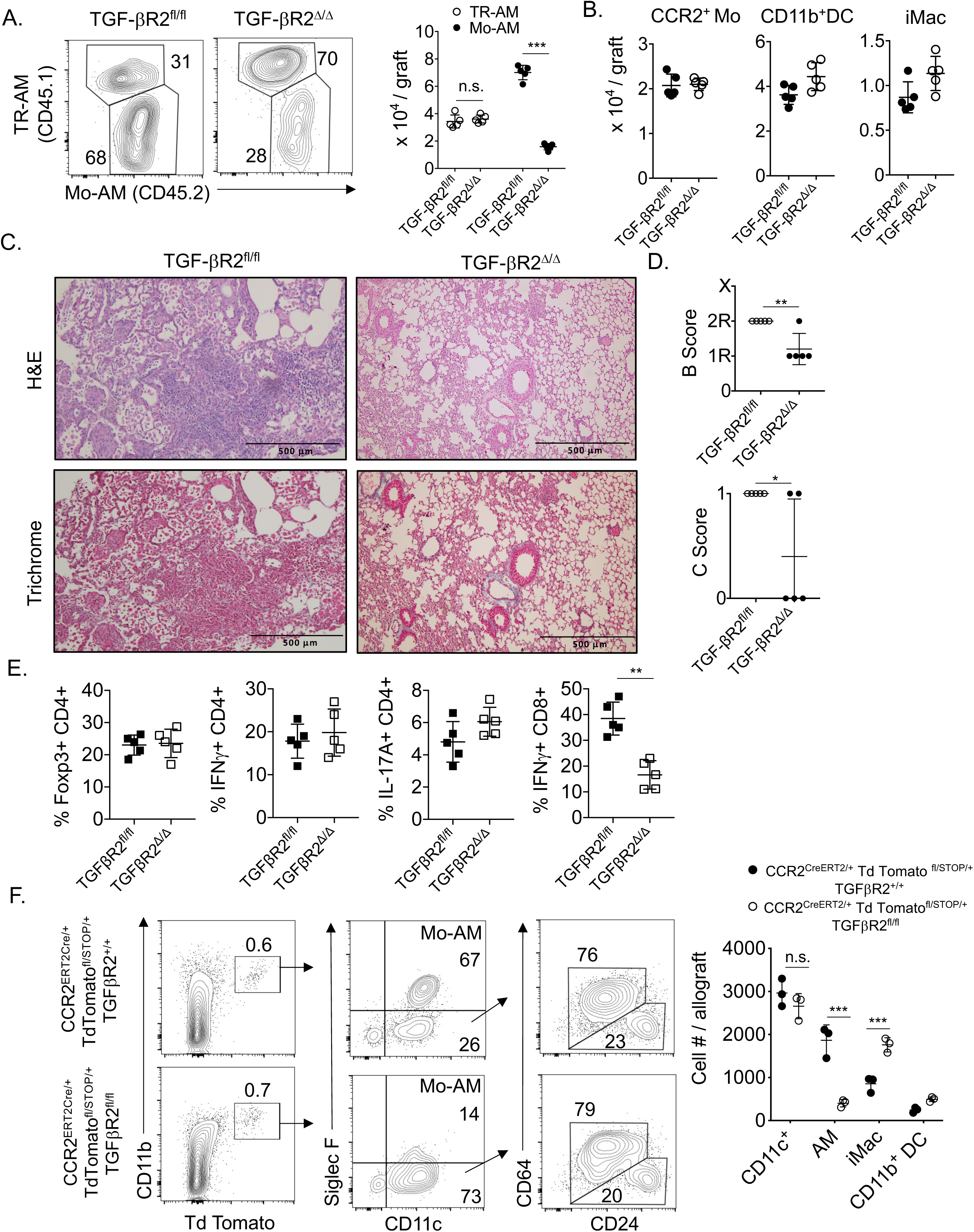
CCR2^+^ monocyte differentiation into Mo-AM requires TGF-β leading to BOS. TGF-βR2^fl/fl^ and TGF-βR2^Δ/Δ^ recipients of 3T-FVB allografts received tamoxifen i.p. every other day for 10 days, rested for 5 days, and then ingested DOX for 2 days. Eight days later, allograft recipients were analyzed for intragraft inflammation **(A)** as shown by representative FACS plots (N=5/group) of the relative percent abundance of Mo-AM and TR-AM with cell counts, **(B)** CCR2^+^ monocyte (Mo), CD11c^+^ DC and iMac cell counts, (**C**) representative H&E and trichrome staining (N=5/group), (**D**) airway inflammation and lesion grading and (**E**) intragraft T cell activation. (**F**) 3 x 10^6^ FACS purified CCR2^+^ bone-marrow monocytes were isolated from indicated Td Tomato reporter mice that received tamoxifen as in (A) and injected into POD 7 3T-FVB recipients undergoing BOS pathogenesis. On POD 16 allograft tissue were quantified for Td Tomato^+^ Mo-AM, CD11b^+^ DC and iMac as shown by representative FACS plots (N=3/group) and cell counts. Bars represent means ± S.D where *p < 0.05; **p < 0.01, *** p < 0.001.

### Mo-AM promote T_RM_ activation and expansion

We have previously demonstrated that donor antigen-primed effector CD8^+^ T cells prevent club cell proliferation and that CD8^+^ T cells are critical mediators of BOS development (*6*). In light of the correlation between Mo-AM and IFN-γ^+^ CD8^+^ T cell accumulation in allografts with BOS we next set out to analyze the expression of surface molecules on Mo-AM that control effector CD8^+^ T cell activation (**Fig. 7A**). Consistent with our previous observations that lung allograft-infiltrating CCR2^+^ monocyte-derived cells express donor-derived MHC molecules, examination of Mo-AM from 3T-FVB transplants revealed the acquisition of the donor-derived MHC Class I molecule H-2K^q^ (*24*) . However, unlike TR-AM, Mo-AM lacked expression of the checkpoint inhibitory molecule PD-L1 and had higher levels of costimulatory ligands CD80 and CD86. We next analyzed patterns of PD-1 in the CD8^+^ T cell compartment of 3T-FVB lung transplants of TGF-βR2^Δ/Δ^, TGF-βR2^fl/fl^ and ECP-treated recipients (**Fig. 7B, Fig. S7A**). In TGF-βR2^fl/fl^ recipients, approximately a third of allograft-resident CD8^+^ T cells co-expressed PD-1 and the integrin CD49a (*52*). In contrast, allografts of TGF-βR2^Δ/Δ^ and ECP-treated recipients had significantly fewer PD-1^+^ CD49a^+^ CD8^+^ T cells. Despite differences in abundance, the PD-1^+^ CD49a^+^ CD8^+^ T cell compartment in all three allograft recipients exhibited a similar T_RM_ phenotype (*53*) (**Fig. 7C**). Nearly all PD-1^+^ CD49a^+^ CD8^+^ T cells were CD44^+^, but lacked expression of CD62L, CCR7 and the killer-like receptor G1 (KLRG1), indicating they were not central memory or short-lived effector cells (*54, 55*). Additionally, PD-1^+^ CD49a^+^ CD8^+^ T cells did not express additional checkpoint inhibitory molecules such as TIM-3 and LAG-3 indicating they were not exhausted memory cells (**Fig. S7B**)(*56*). Moreover, we detected a similar T_RM_ phenotype and abundance in 2T-FVB allografts demonstrating that these cells exist in accepted lung transplants prior to the development of BOS (**Fig. S7C**). However, PD-1^+^ CD49a^+^ CD8^+^ T cells in allografts of TGF-βR2^fl/fl^ recipients expressed moderate levels of the T_RM_ marker CD103 and high levels of Gzmb and Blimp-1.

**Figure 7:**
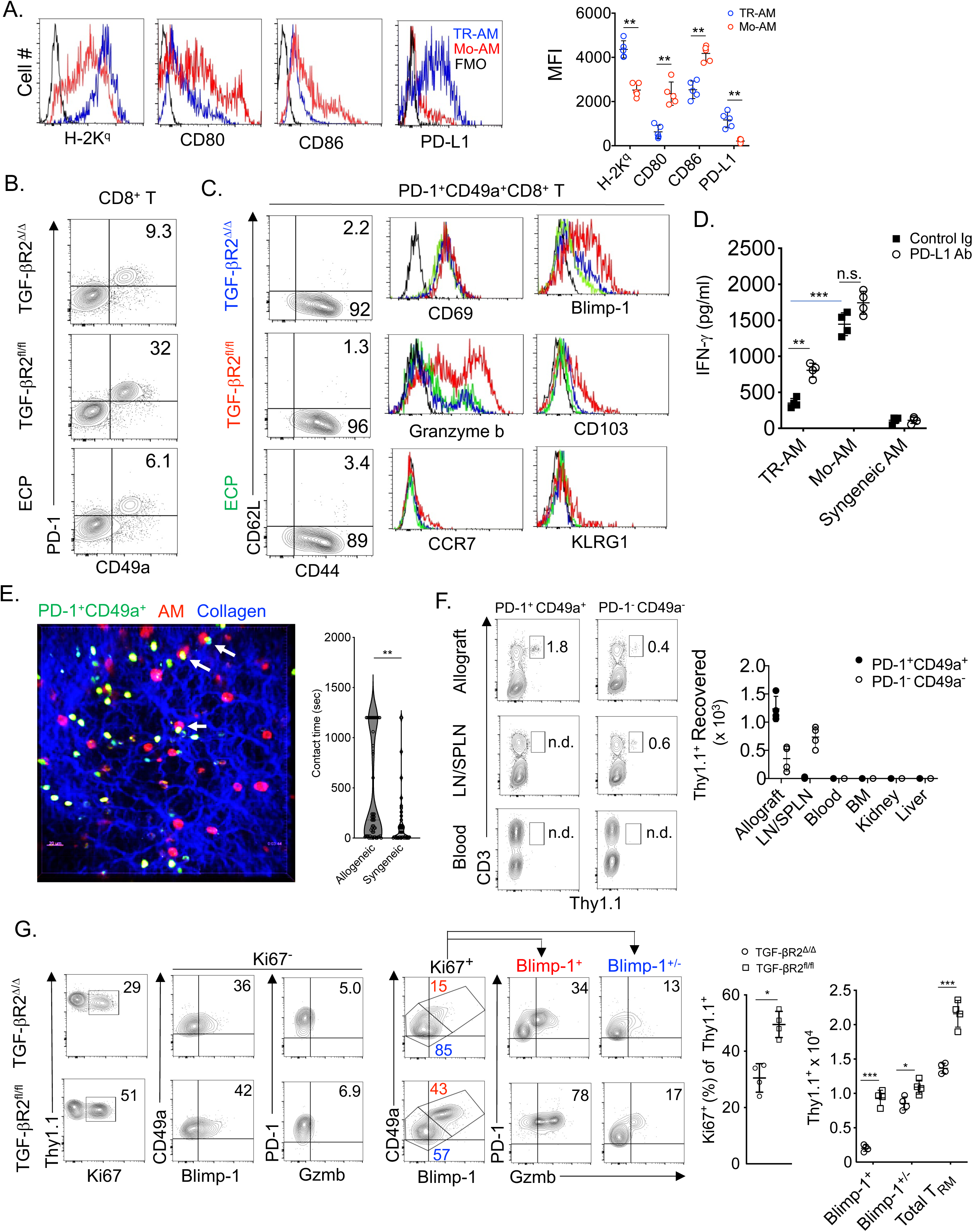
Mo-AM generation promotes T_RM_ activation and expansion. **(A)** Representative FACS plots and absolute MFI of TR-AM and Mo-AM MHC I H-2K^q^, CD80, 86 and PD-L1 expression levels with FMO (fluorescence minus one control). (**B-C**) Representative N=4/group) FACS plots and histograms of T_RM_ expression patterns with FMO (black lines). (**D**) FACS -sorted 3T-FVB TR-AM, Mo-AM, and B6 AM were cultured with FACS-sorted 3T-FVB allograft PD-1^+^ CD49a^+^ CD8^+^ T cells with 10 μg/ml control rat Ig or PD-L1 neutralizing Abs and then assessed for IFN-γ production by ELISA 72 hours later. (**E**) FACS-sorted, CFSE- labeled 3T-FVB allograft PD-1^+^ CD49a^+^ CD8^+^ T cells (Green) were intratracheally administered to FVB (Allogeneic) or B6 (Syngeneic) lung transplants of B6 recipients and imaged by 2- photon intravital microscopy 18 hours later following the administration of Siglec F antibodies to identify AM (Red). Representative image of a transplanted FVB lung (N=4) where arrows denote long-lasting contacts between T_RM_ and AM. Right panel is violin plot of individual AM- T_RM_ contact times for 1200 sec imaging time frames. (F) 2T-FVB allografts of B6 Thy 1.1+ recipients were FACS-sorted for PD-1^+^ CD49a^+^ and PD-1^-^ CD49a^-^ CD8^+^ T cells and intratracheally delivered into B6 Thy 1.2^+^ recipients of 2T-FVB allografts and euthanized one month later. Data shown are representative FACS plots (N=4/group) of Thy1.1^+^ cell percent abundance and cell count in indicated tissues. (G) 2T-FVB allograft FACS-sorted Thy1.1 PD-1^+^ CD49a^+^ CD8^+^ T cells were intratracheally administered into tamoxifen-treated TGF-βR2^fl/fl^ and TGF-βR2^Δ/Δ^ recipients of 3T-FVB allografts three days after DOX ingestion. 72 hours later recipients were euthanized. Data shown are representative FACS plots (N=4/group) of allograft percent abundance and cell counts. Bars represent means ± S.D where *p < 0.05; **p < 0.01, *** p < 0.001, n.s.; not significant.

Viral peptide-and alloantigen-specific T_RM_ can become reactivated upon cognate antigen encounter (*28, 57*). We next isolated PD-1^+^ CD49a^+^ CD8^+^ T cells from 3T-FVB allografts with BOS and measured IFN-γ expression in response to stimulation with 3T-FVB allograft-derived Mo-AM and TR-AM (**Fig. 9D**). TR-AM were poor at eliciting IFN-γ production when compared to Mo-AM. However, the addition of anti-PD-L1 antibodies to TR-AM, but not Mo- AM co-cultures significantly increased IFN-γ responses. Although these data indicated that TR- AM and Mo-AM differentially regulate T_RM_ activation responses through PD-L1 expression, it remained unclear if these cells directly interact with AM within lung transplants. To answer this question, we utilized intravital two-photon microscopy to assess contact times between PD-1^+^ CD49a^+^ CD8^+^ T cells and AM (**Fig. 7E, Movie S1, S2**). CFSE-labeled PD-1^+^ CD49a^+^ CD8^+^ T cells isolated from 3T-FVB allografts were intratracheally delivered into FVB lung (allogeneic) or control syngeneic B6 lung recipients. One day later, lung recipients also received Siglec-F fluorescently-labeled antibodies to identify AM. Relative to syngeneic B6 lung transplants, significantly prolonged interactions were observed between AM and PD-1^+^ CD49a^+^ CD8^+^ T cells in FVB allografts, which is indicative of donor-antigen recognition (*58, 59*). A canonical property of T_RM_ is their inability to exit from barrier organs to recirculate in the periphery (*57*). To determine if this was the case for PD-1^+^ CD49a^+^ CD8^+^ T cells, we isolated Thy1.1^+^ PD-1^+^ CD49a^+^ CD8^+^ T cells from 2T FVB allografts of Thy1.1^+^ B6 recipients and intratracheally delivered these cells into 2T FVB allografts of Thy1.2^+^ B6 recipients (**Fig. 7F**). One month later, we could detect Thy1.1^+^ cells in lung allograft tissue, but not in secondary lymphoid organs, peripheral blood, bone marrow, liver or kidney. In contrast, 2T-FVB allograft-derived Thy1.1^+^ PD-1^-^ CD49a^-^ CD8^+^ T cells were detected in secondary lymphoid organs indicating that intratracheally administered CD8^+^ T cells can exit lung allografts. Therefore, lung allograft PD- 1^+^ CD49a^+^ CD8^+^ T cells are phenotypically and functionally consistent with T_RM_ and herein we will refer to these cells as T_RM_.

Recent work in models of cutaneous viral infection has indicated that T_RM_ expand from a local pre-existing population of T_RM,_ but whether this is true for pulmonary T_RM_ is less understood (*60*). Because we noted that T_RM_ are more abundant in BOS compared to accepted allografts, we asked if Mo-AM generation during BOS development drives the local expansion of these cells from pre-existing intragraft pools. Therefore, we adoptively transferred Thy1.1^+^ T_RM_ from accepted 2T FVB allografts into 3T FVB allografts airways of TGF-βR2^fl/fl^ and TGF- βR2^Δ/Δ^ Thy1.2^+^ recipients following tamoxifen treatment and bronchiolar injury (**Fig. 7G**). Five days later, intragraft Thy1.1^+^ T_RM_ were analyzed for proliferation and accumulation. In allografts of TGF-βR2^fl/fl^ recipients, Thy1.1^+^ T_RM_ cells proliferated and accumulated at significantly higher levels when compared to TGF-βR2^Δ/Δ^ recipients. Analysis of the proliferating Thy1.1^+^ T_RM_ compartment of TGF-βR2^fl/fl^ allografts recipients revealed high numbers of PD-1^+^ Blimp-1^+^ CD8^+^ T cells that expressed elevated Gzmb, which was similar in phenotype to the native T_RM_ phenotype detected in these allografts. Additionally, both allografts contained proliferating PD-1^+^ Blimp-1^+/-^ Gzmb^+/-^ CD8^+^ T cells, a phenotype that resembled the native T_RM_ allograft compartment observed in TGF-βR2^Δ/Δ^ and ECP-treated recipients. Finally, we could detect clusters of Gzmb^+^ CD49a^+^ CD8^+^ cells in explanted lung transplant tissue from BOS patients that were not present in stable recipients that do not have evidence of rejection (**Fig. S8**). Collectively, our observations indicate that the Mo-AM generation during BOS pathogenesis drives the activation and expansion of T_RM_.

### T_RM_ Gzmb expression induces airway epithelial apoptosis and promotes BOS

Gzmb induces mitochondrial stress leading to apoptosis and has been reported to be elevated in the BALF of BOS subjects (*61, 62*). The finding of high Gzmb expression in TRM from allografts with BOS indicated the potential to promote airway epithelial cell cytotoxicity. We isolated T_RM_ from 3T-FVB allografts with BOS for co-culture with lung epithelial cells from FVB mice and measured changes in mitochondrial membrane potential, mitochondrial ROS production and DNA fragmentation in the presence or absence of the Gzmb inhibitor Serpin A3N (*63*) (**Fig. 8A, B**). T_RM_ induced rapid mitochondrial stress as evidenced by mitochondrial membrane depolarization and elevated superoxide production. Additionally, DNA fragmentation, an indicator of late-stage apoptosis, was more than fivefold greater relative to control naïve B6 CD8^+^ T cell co-cultures. In contrast, T_RM_-induced mitochondrial stress and DNA fragmentation could be significantly prevented by pre-treatment with Serpin A3N. These data indicate that BOS allograft-derived T_RM_ induce airway injury through Gzmb expression.

**Figure 8:**
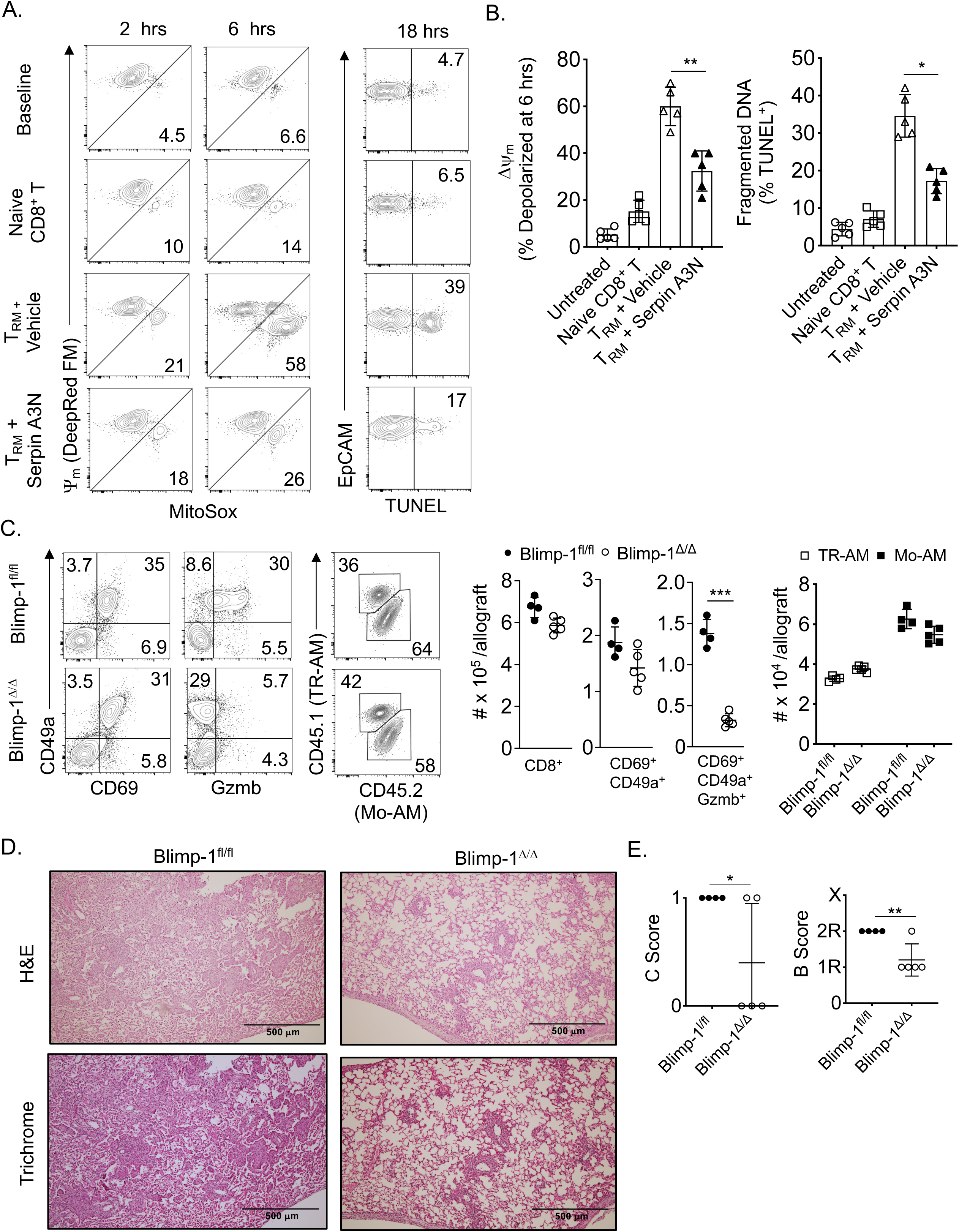
Gzmb^+^ T_RM_ promote airway epithelial cell apoptosis and BOS through Blimp-1. FVB lung epithelial cells were co-cultured in a 1: 2 EpCAM ^+^ cell to CD8^+^ T cell ratio for up to 18 hours with or without (saline vehicle) Serpin A3N pretreatment (25 nM) and assessed for survival mitochondrial charge (MitoTracker Deep Red FM), superoxide production (MitoSOX) and DNA fragmentation (TUNEL). Data shown are representative (**A**) FACS plots (N=5/condition) and (**B**) histograms of 6 hr mitochondrial depolarization and TUNEL activity. Blimp-1^fl/fl^ and Blimp-1^Δ^*^/^*^Δ^ recipients of 3T-FVB allografts were analyzed for intragraft inflammation as shown by (**C**) representative FACS plots of percent abundance (N=5/group) and cell counts, (**D**) representative H&E and Trichrome staining (N=5/group) and (**E**) airway inflammation and lesion grading. Bars represent means ± S.D where *p < 0.05; **p < 0.01.

Blimp-1 has been shown to drive Gzmb expression in mouse T_RM_ (*32*). The observation of co-expression of Blimp-1 with high amounts of Gzmb in T_RM_ from allografts with BOS raised the possibility that it plays a role in promoting rejection. We next used *CD8a*^Cre^ *Pdrm1*^fl/fl^ (Blimp-1^Δ^*^/^*^Δ^) mice as recipients for 3T-FVB lungs and assessed intragraft inflammation and BOS severity. When compared to *Pdrm1*^fl/fl^ (Blimp-1^fl/fl^) recipients, we observed similar numbers of total intragraft CD8^+^ T cells and T_RM,_ with little effect on CD69 and CD49a expression (**Fig. 8C, Fig. S9**). However, Blimp-1^Δ^*^/^*^Δ^ T_RM_ were largely devoid of Gzmb expression and their allografts were markedly protected from severe BOS despite maintaining high numbers of Mo-AM (**Fig. 8D, E**).

## DISCUSSION

Devising therapies to prevent or treat chronic rejection is one of the major challenges in the transplantation field. Advances in this field are largely hampered by our incomplete understanding of the contributory immune mechanisms that could be potentially targeted. Given some encouraging reports of ECP treatment in BOS patients, we reasoned that we could gain insight into the underlying mechanisms of BOS through modeling this therapy in mouse orthotopic lung transplants. Consistent with observations in ECP-treated subjects with BOS (*12*), we noted a significant reduction in lung transplant antigen-specific responses in our model. We also made the novel and clinically relevant observation that ECP ameliorates OB lesion severity (*11*).

Most of the investigative focus into TGF-β-mediated fibrogenesis has led to the elucidation of mechanisms that control extracellular matrix remodeling, epithelial to mesenchymal transition and fibrogenesis. Although these pathways are involved in BOS pathogenesis (*64*), the additional requirement for leukocyte-dependent recognition of alloantigens mechanistically distinguishes BOS from other pulmonary fibrotic diseases. We were initially surprised by reports of ECP’s effectiveness in BOS patients, given that previous studies have demonstrated that ECP stimulates TGF-β protein expression along with the expansion of Foxp3^+^ CD4^+^ T cells (*13*). However, we did not find that ECP increased TGF-β protein expression or Foxp3^+^ CD4^+^ T cells, which suggests that different mechanisms drive ECP-mediated immunoregulatory effects in lung transplant recipients. Surprisingly, we discovered that ECP inhibits lung allograft airway TGF-β activity by inducing DCN expression by AM. We contend that local TGF-β activity is the principal therapeutic target of ECP in lung transplants, which is also supported by our finding that intratracheal administration of TGF-β- neutralizing antibodies also inhibited BOS.

Non-invasive approaches to detect early BOS development have yet to be developed due to a lack of knowledge of the underlying immunological mechanisms that lead to OB. Using CCR2^+^ probe-based micro-PET imaging, we detected a large increase in CCR2^+^ intragraft activity in lung recipients with BOS that was largely reversed by ECP treatment. It should be noted that the detection of intragraft CCR2 activity could be due to recruitment or development of CCR2^+^ macrophages. However, since large numbers of CCR2^+^ monocytes can be detected in lung allografts with BOS and BOS development is inhibited in CCR2-monocyte depleter lung recipients, our results suggest that increased PET activity is primarily due to the infiltration of CCR2^+^ monocytes.

A recent study employing single cell RNA sequencing revealed that the majority of AM in human lung allografts are derived from the recipient (*65*). These observations led us to consider the relevance of Mo-AM accumulation during BOS development. Notably, we observed a high Mo-AM to TR-AM ratio in BOS allografts that was driven by a combination of CCR2^+^ monocyte differentiation and a reduction in TR-AM numbers following bronchiolar injury. The reasons for the loss of TR-AM are not clear, but it was not preventable by ECP or anti-TGF-β antibody treatment. Therefore, their loss could be potentially explained by their programmed cell death following airway inflammation (*66*). We recognized that targeting CCR2-monocyte-mediated Mo-AM depletion could also result in the defective generation of other CCR2^+^ monocyte-derived cells that may contribute to BOS, such as iMacs and CD11b^+^ DCs (*24*). Following total body irradiation, lysosomal M-mediated expression of TGF-βR2 has been reported required for AM reconstitution raising the possibility that CCR2^+^ monocytes give rise to Mo-AM (*23*). We conducted monocyte fate studies with reporter mice in which TGF-βR2 deletion and Td tomato expression are both under the inducible control of CCR2 cre recombinase. In recognition of the leakiness of the Td-tomato ‘flox on’constructs (*67*), we studied the differentiation of purified CCR2^+^ monocytes following adoptive transfer into lung recipients undergoing BOS pathogenesis. We found that CCR2^+^ monocyte differentiation into Mo-AM is substantially dependent on TGF-β signaling during BOS pathogenesis. However, for iMacs, we observed comparatively less generation from wild type monocytes indicating TGF-β signaling may retard their development. Additionally, it is important to note that we could detect small numbers of Mo-AM in tolerant allografts, indicating that Mo-AM generation is not sufficient to promote BOS. Interestingly, reports exist that clodronate-mediated TR-AM depletion prior to bleomycin treatment does not worsen pulmonary fibrosis (*21*). We also observed that clodronate-mediated donor TR-AM depletion and subsequent Mo-AM reconstitution does not spontaneously induce or increase the severity of BOS. Thus, our data point to the requirement for airway inflammation to trigger Mo-AM-dependent BOS development.

We found that PD-1 expression on intragraft CD8^+^ T cells largely marked the T_RM_ compartment irrespective of tolerance status. Recent work in a mouse model of acute influenza infection has demonstrated that MHC Class I, CD80 and CD86 are all required to maintain PD- 1^+^ T_RM_ (*68*). Interestingly, T_RM_ in this setting were found to be exhausted as PD-L1 blockade was required to clear secondary infection at the cost of developing fibrotic sequelae. In contrast, repeated PD-1 antibody blockade in a mouse kidney allograft model highly enriched for PD-1^+^ T_RM_ failed to exacerbate chronic rejection (*57*). Similar to our observations in the current study, these investigators did not find evidence of T_RM_ exhaustion. Lung allograft T_RM_ did not co-express additional exhaustion markers and robustly recalled IFN-γ expression upon donor-antigen challenge by Mo-AM. In contrast, T_RM_ reactivation by TR-AM was poor largely due to PD-L1 expression. These data therefore suggest that TR-AM may limit alloimmune responses. Alternatively, TR-AM could contribute to T_RM_ development as a recent report demonstrated that AM depletion prevents T_RM_ differentiation in a murine influenza infection model (*29*). Overcoming PD-1-mediated inhibition of CD8^+^ T cell memory responses requires engagement CD80 and CD86 (*69*), two co-stimulatory ligands expressed on both AM subsets. Our group has previously demonstrated that PD-1 expression on CD8^+^ T cells is required for costimulatory blockade-mediated induction of lung transplant acceptance as well as for prolonged interactions with recipient-derived intragraft CD11c^+^ cells (*70*). In this study, we detected prolonged interactions between allograft T_RM_ and AM that were donor-antigen dependent. Future intravital studies will be needed to assess whether T_RM_ interactions with TR-AM are also dependent on PD-1/PD-L1 engagement.

Unique to allografts with BOS is the accumulation of Gzmb^hi^ Blimp-1^+^ T_RM_. Using an airway epithelial cell co-culture system, we observed rapid and potent Gzmb-dependent apoptotic activity by BOS allograft T_RM_ that is suggestive of potent pathogenic activity. We also investigated the origins and requirements of the Gzmb^hi^ Blimp-1^+^ T_RM_ subset. Previous work in virally infected mice has demonstrated that skin T_RM_ can maintain themselves locally from a pool of pre-existing T_RM_ (*60, 71*). Our finding that Blimp-1^+^ Gzmb^hi^ T_RM_ can be generated from T_RM_ isolated from tolerant allografts is in line with these previous reports and suggests that specifically targeting intragraft T_RM_ may be a viable strategy to prevent BOS. However, our studies do not rule out a possible contribution from peripheral T_RM_ precursors. Insight into the in vivo antigen presenting cell requirements for Blimp-1^+^ Gzmb^hi^ T_RM_ generation was gained by observations of sharply lower numbers of these cells in TGF-βR2^Δ^*^/^*^Δ^ recipients, which were devoid of Mo-AM, but not other CD11c^+^ antigen presenting cells. When these data are considered in conjunction with ability of Mo-AM to induce T_RM_ IFN-γ expression, our results support the notion that Mo-AM-mediated antigen presentation is a key promoter of T_RM_ activation and expansion.

Blimp-1 expression promotes T_RM_ Gzmb, but appears dispensable for CD49a or CD69 expression (*32, 72, 73*). This relationship allowed us to analyze the effects of T_RM_-mediated Gzmb expression on allograft tolerance. Blimp-1 ablation within the CD8^+^ T cell compartment led to a drastic reduction of T_RM_ Gzmb expression and inhibited BOS development. Nevertheless, it remains plausible that protection from BOS could be due to defective T_RM_ or effector CD8^+^ T cell development unrelated to Gzmb expression. A recent report has shown that Blimp-1 is required for lung CD103^+^ T_RM_ development (*72*). We observed mild CD103 expression in BOS allograft T_RM_. When compared to CD49a and CD69, which play well-established roles in survival, trafficking and retention, the role of CD103 is less clear, but could act to promote epithelial adherence (*74*). Nevertheless, recent work has demonstrated that CD49a is a critical marker of T_RM_ cytolytic activity irrespective of CD103 expression (*75*) . Moreover, considering our observations of CD8^+^ CD49a^+^ Gzmb^+^ cells in BOS subjects and several previous outcome studies demonstrating an association between airway Gzmb^+^CD8^+^ T cell accumulation and risk for BOS, our data collectively point to T_RM_ Gzmb expression as an important contributor to OB lesion development (*76*). Finally, we observed that Blimp- 1^Δ^*^/^*^Δ^ allograft recipients generated high numbers of Mo-AM providing further evidence that this AM subset drives BOS by promoting T_RM_ expansion and cytotoxicity.

In conclusion, our work reveals the existence of an inducible functional plasticity within the AM compartment that can be harnessed to lower TGF-β bioavailability and prevent BOS. These findings also extend the notion that AM have the capacity to alter their local environment (*77*). Our results also uncover TGF-β-mediated Mo-AM development as a regulator of adaptive immunity with implications beyond the transplantation field.

## MATERIALS AND METHODS

### Mice and Orthotopic Lung Transplantation

C57BL/6 (B6), FVB, Thy1.1, Lyz^Cre^, CD8^Cre^, Prdm1^fl/fl^ and Tgfbr2^fl/fl^ mice were all purchased from Jackson Labs. CCR2^DTR^, Dcn^fl/fl^ and CCR2^CreERT2^ mice were generous of gifts from Prof. Eric G. Pamer of the University of Chicago, Prof. David E. Birk of the University of South Florida and Prof. Dr. Burkhard Becher of the University of Zurich, Switzerland, respectively. 2T-FVB and 3T-FVB donor mice and mouse left orthotopic lung transplantation procedures have been previously described by our group (*6*). To induce allograft acceptance, recipients received i.p. 250 μg of CD40L Abs (MR1) on POD 0 and 200 μg of mouse recombinant CTLA4Ig on POD 2 (*78*). Club cell injury is triggered by DOX ingestion via food (625 mg/kg chow; ENVIGO) and water (2 mg/ml, Sigma) for 2 to 2.5 days. Tamoxifen (Sigma-Millipore) was dissolved in Mazola corn oil and injected i.p. five times at 0.25 mg/g body weight every other day and then rested 5 days prior to DOX ingestion. Diptheria toxin (Sigma-Millipore) was dissolved in PBS and injected i.p. one day prior to DOX ingestion at 10 ng/g body weight.

### ECP

B6 (syngeneic) leukocytes were isolated from splenocytes by centrifugation through a density separation medium (Lympholyte-M, Cedarlane) to eliminate dead cells, debris and erythrocytes. Remaining erythrocytes were removed by ACK lysing buffer, and leukocytes were resuspended at 5 x 10^6^ cells/ml in complete medium with 8-methoxypsoralen (8-MOP, 200 ng/mL), incubated in the dark for 30 mins at 25 °C and irradiated at 2 J/cm^2^ UVA in a ECP irradiator box (Johnson & Johnson). Following DOX ingestion, recipients received three i.v. doses of 10^7^ ECP-treated leukocytes in 100 μl normal saline spaced at three-day intervals.

### Histological staining and analysis

Harvested grafts were formaldehyde-fixed, paraffin-embedded and stained with hematoxylin-eosin or Masson’s trichrome stain. Lung transplant histology was graded by a blinded pathologist using the 2007 revision of the 1996 working formulation for the standardization of nomenclature in the diagnosis of lung rejection. For immunohistochemical analysis of bronchiolar epithelium paraffin sections were first blocked with 5% goat serum and 2% fish gelatin (Both from Sigma-Aldrich) at 25 ^0^C for 45 mins. Sections were then stained with 1:500 polyclonal rabbit anti-mouse/rat CCSP (Seven Hills Bioreagents) and mouse anti-Acetylated Tubulin, 1:5000 (6-11B-1, Sigma-Aldrich) overnight at 4 ^0^C. For immunofluorescent visualization, we used goat anti-mouse Alexa Fluor 488-labeled secondary antibody, 1:1000 (Thermo Fisher) and goat anti-rabbit Alexa Fluor 555-labeled secondary antibody, 1:1000 (Cell Signaling Technology). Mouse-anti human CD8α (C8/114B), rat anti-human (16G6) and rabbit anti-human polyclonal CD49a were all purchased from ThermoFisher. For collagen measurements 10 mg of allograft tissue/sample was analyzed with a Hydroxyproline Assay Kit (Sigma) in accordance with manufacturers recommendations.

### Flow Cytometric Analysis and Antigen Recall Assays

Lung tissue was minced and digested in a RPMI 1640 solution with Type 2 collagenase (0.5 mg/mL) (Worthington Biochemical) and 5 units/mL DNAse (Sigma) for 90 min at 37 ^0^C and then filtered through a 70-um cell strainer (ThermoFisher) and treated with ACK lysing buffer (Worthington Biochemical). Live cell discrimination was conducted with the Zombie (Biolegend) fixable dye. Cell surface staining was conducted with the following Abs; CD45 (30-F11; eBioscience), CD45.2 (104; Biolegend), CD90.2 (53-2.1; eBioscience), CD4 (clone RM4-5; eBioscience), CD8α (53-6.7; eBioscience), CD31 (390; Biolegend), CD34 (HM34; Biolegend) and CD326 (G8.8; Biolegend). Staining for Foxp3 (FJK-16s, eBioscience) and Ki-67 (16A8; Biolegend) and CCSP (Seven Hills Bioreagents) was conducted with intranuclear Transcription Factor Staining Buffer Kit (Invitrogen) in accordance with manufacturers recommendations. For IFN-γ and IL-17A expression cells were first stimulated with1 μM ionomycin (Sigma) and 20 ng/ml PMA (Sigma) for 3.5 hrs with 2 μM Golgi Plug (BD Biosciences) added for last 3 hrs of stimulation and stained with IFN-γ (XMG1.2; eBioscience) and IL-17A (TC11-18H10.1; Biolegend) using a Cytofix/Cytoperm kit (BD Biosciences) in accordance with manufacturers recommendations. For antigen specificity measurements, T cells were fractionated by positive selection using CD4^+^ or CD8^+^ immunomagnetic beads (Miltenyi) from allograft cell suspensions and co-cultured in a 3:1 ratio with irradiated T cell-depleted FVB or B6 cell splenocytes for 96 hours pulsed with 0.5 µg/ml K-α1 tubulin and Collagen V (obtained from T. Mohannakumar, St. Joseph’s Hospital, Phoenix AZ). IFN-γ and IL-17A were measured with uncoated ELISA kits from Invitrogen in accordance with manufacturer’s recommendations.

### Semi-quantitative RT PCR

AM were extracted for RNA with RNAeasy kits (Qiagen) and reverse transcribed with the high capacity cDNA reverse transcription kit (Thermo Fisher). Transcripts were semi-quantified with Taqman Gene Expression Assays for indicated genes (Thermo Fisher) where transcript levels were normalized against the macrophage housekeeping gene *Sxt5a* (*79*).

### TGF-β measurements

TGF-beta isoforms were measured with the Bio-Plex Pro TGF-β assays (Bio-Rad) in accordance with manufactures directions. To measure TGF-β activity, NIH/3T3 SMAD2/3-luciferase reporter cell line (Signosis) was co-cultured with BALF, extracted with RPMI 1640 (3:1 v/v ratio) or platelet-free plasma in a 1:10 (v/v) ratio for 16 hrs. Luciferase activity was measured using BioTek Synergy/HTX Multi-mode reader.

### Cell culture

Mo-AM and TR-AM were isolated by FACS-sort as CD45.2^+^ or CD45.1^+^ CD11c^+^ Siglec F^+^CD11b^+^ CD64^+^ cells from either 3T-, 2T-FVB allograft recipients or B6 mice and seeded into 96 well round bottom plates at 5.0 to 7.5 x 10^4^ per well and co-cultured with PD-1^+^CD49a^+^CD8^+^ T cells FACS-sorted from either 2T or 3T FVB allograft recipients in a 1:1 ratio in the presence of 10 μg/ml anti-mouse PD-L1 Abs (BE0361; Bio-X-Cell) or control Rat IgG for 72 hours. Cultures were stimulated with 20 ng/ml PMA for 3 hours and assessed with a mouse IFN-γ ELISA kit (Sigma-Millipore). For airway epithelial cell culture, FVB lung tissue cell isolates were prepared as described for FACS preparation and incubated with biotin-conjugated Abs (all from eBioscience) specific for CD45.1 (clone A20), CD34 (clone RAM34), CD31 (clone MEC13.3), CD90.1 (clone HIS51), and CD15 (clone mc-480), washed, and then labeled with anti-biotin MicroBeads (Miltenyi Biotec) for negative selection on LS columns (Miltenyi Biotec). Remaining cells were then incubated with biotin-conjugated CD326 Abs (clone caa7-9G8, Miltenyi Biotec), washed, and then labeled anti-biotin MicroBeads for MS column (Miltenyi Biotec) mediated-positive selection. Enriched club cell fractions were resuspended in MTEC/Plus medium and seeded at 3.0 ×10^4^ cell per well in flat bottom 96 well tissue culture plates (Thermo Fisher) coated with 50 μg/ml type I rat tail collagen (Becton-Dickinson). 3T-FVB allograft T_RM_ were added at epithelial cells in a 1:1 ratio and Mitotracker DeepRed FM and MitoSOX (Both from Thermofisher) were added at 1 μM to cultures 30 mins prior to removal for FACS analysis. DNA fragmentation was measured by TUNEL FACS-based assay kit (Abcam) in accordance manufacturers recommendation.

### PET Imaging

0–60 min dynamic PET/CT scan was performed following injection of ^64^Cu-DOTA-ECL1i (100 μCi in 100 μL saline) with microPET Focus 220 (Siemens, Malvern, PA) or Inveon PET/CT system (Siemens, Malvern, PA). The PET images were reconstructed with the maximum a posteriori algorithm and analyzed by Inveon Research Workplace. The organ uptake was calculated as percent injected dose per gram (%ID/g) of tissue in three-dimensional regions of interest without correction for partial volume effect. Reagents, synthesis and characterization of all compounds have been previously described by our group (*80*).

### 2-photon intravital imaging

Chest wall exposure was conducted between the 3^rd^ and 7th ribs and a cover glass slide was adhered to the lung allograft using tissue glue (VetBond) applied in a gentle manner as not to disturb blood flow. AM were imaged PE-Siglec F (2 µg, clone S17007L; BioLegend) administered intravenously 30 mins after engraftment. FACS-sorted PD-1^+^ CD49a^+^ CD69^+^ CD8^+^ T cells isolated from e3T-FVB allografts were labeled with 5 μM CFSE and between 1 and 3 x 10^5^ cells were intratracheally administered one day before imaging. Data was collected by sequential z-sections (24, 2.5 μm each) were acquired in an imaging volume of 200 × 225 × 60 μm^3^. Analyses were performed with Imaris (Bitplane, Zurich, Switzerland). Associations between AM and T_RM_ were defined as physical interactions that lasted greater than 15 secs. For each lung transplant, at least 5 areas were examined up to approximately 50 μm deep. Data shown has been pooled from at least 3 mice per group.

### Statistics

Data are presented as mean ± S.D. The student t test or Mann–Whitney U test was performed by GraphPad Prism version 7.0 (GraphPad Software). P < 0.05 was considered significant.

## Acknowledgements

We would like thank Drs. Mark J. Miller and Seonyoung Kim of the Washington University In Vivo Imaging Core for technical advice.

## Funding

Cystic Fibrosis Foundation (AEG)

The Barnes Jewish Foundation (AEG)

National Instutitutes of Health P01AI116501 (ASK, DK, AEG)

National Instutitutes of Health P41EB025815 (YL, AEG)

National Instutitutes of Health R01HL151685 (YL)

National Instutitutes of Health R35HL145212 (YL)

National Instutitutes of Health R01HL094601 (DK, AEG)

National Instutitutes of Health K08HL148510 (HSK)

The Children’s Discovery Institute PD-FR-2020-867 (HSK)

National Instutitutes of Health K01HL155231 (LTK)

The Harold Amos Medical Faculty Development Program (LTK)

## Authors Contributions

Investigation: ZL, FL, JZ, DZ, GSH, HPL, DS, AP, MC, WL and AEG

Methodology: RH, DB, LKT, HJH, BWW, HSK and AEG

Conceptulization: HJH, YL and AEG

Funding Acquisition: YL, ASK, DK and AEG

Writing: YL, DK and AEG

## Data and materials avialability

All data are available in the main text or the supplementary materials

## Supplementary Materials

**Movie S1. T_RM_ make prolonged associations with allogeneic lung transplant AM.** CFSE- labeled PD-1^+^ CD49a^+^ CD8^+^ T cells isolated from 3T-FVB allografts of B6 recipients and adoptively transferred into FVB lung transplants of B6 recipients. One day later PD-1^+^ CD49a^+^ CD8^+^ T cells (Green) were analyzed for association with AM labeled with Siglec-F Ab conjugated to phycoerythrin (Red). Collagen fibers are represented by the blue color of secondary harmonic fluorescence. Data shown is a representative 20-minute movie from 4 experiments.

**Movie S2. T_RM_ do not make prolonged associations with syngeneic lung transplant AM.** CFSE-labeled PD-1^+^ CD49a^+^ CD8^+^ T cells isolated from 3T-FVB allografts of B6 recipients and adoptively transferred into B6 lung transplants of B6 recipients. One day later PD-1^+^ CD49a^+^ CD8^+^ T cells (Green) were analyzed for association with AM labeled with Siglec-F Ab conjugated to phycoerythrin (Red). Collagen fibers are represented by the blue color of secondary harmonic fluorescence. Data shown is a representative 20-minute movie from 4 experiments.

---

**Fig. S1:**
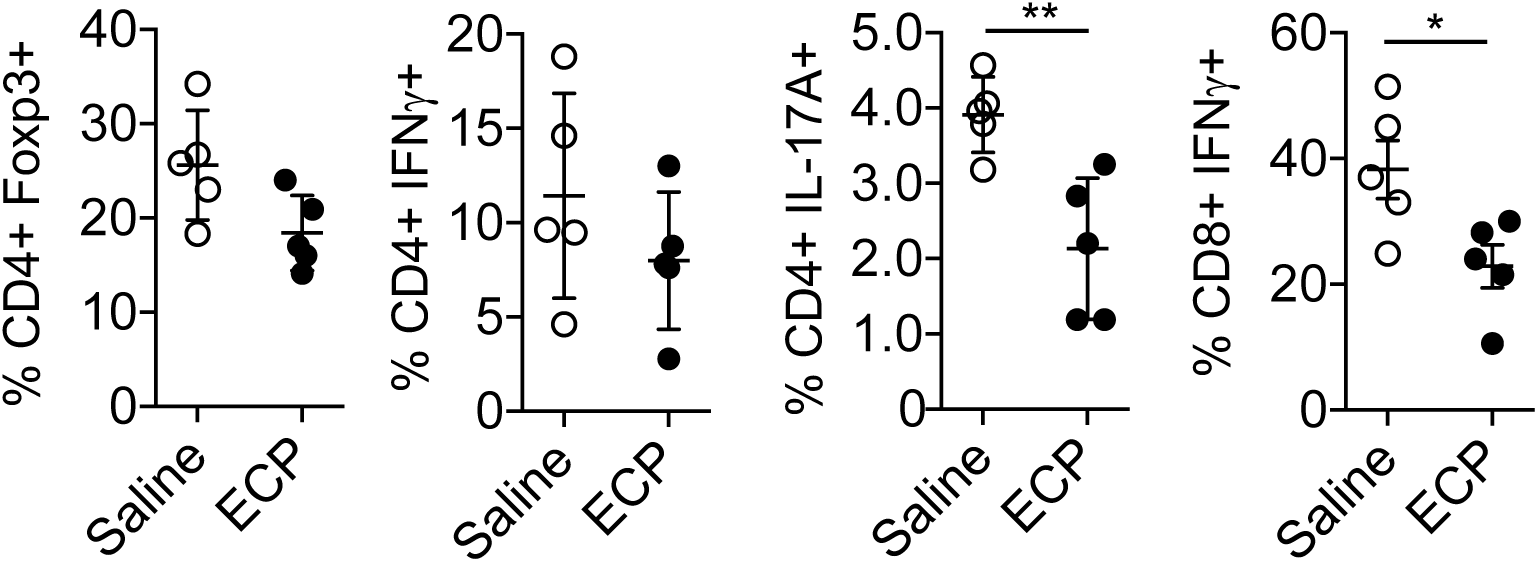
ECP reduces T_h_17 and effector CD8^+^ T cell abundance. Intragraft abundance for indicated T cell lineages. Bars represent means ± S.D where *p< 0.05, **p < 0.01.

**Fig. S2:**
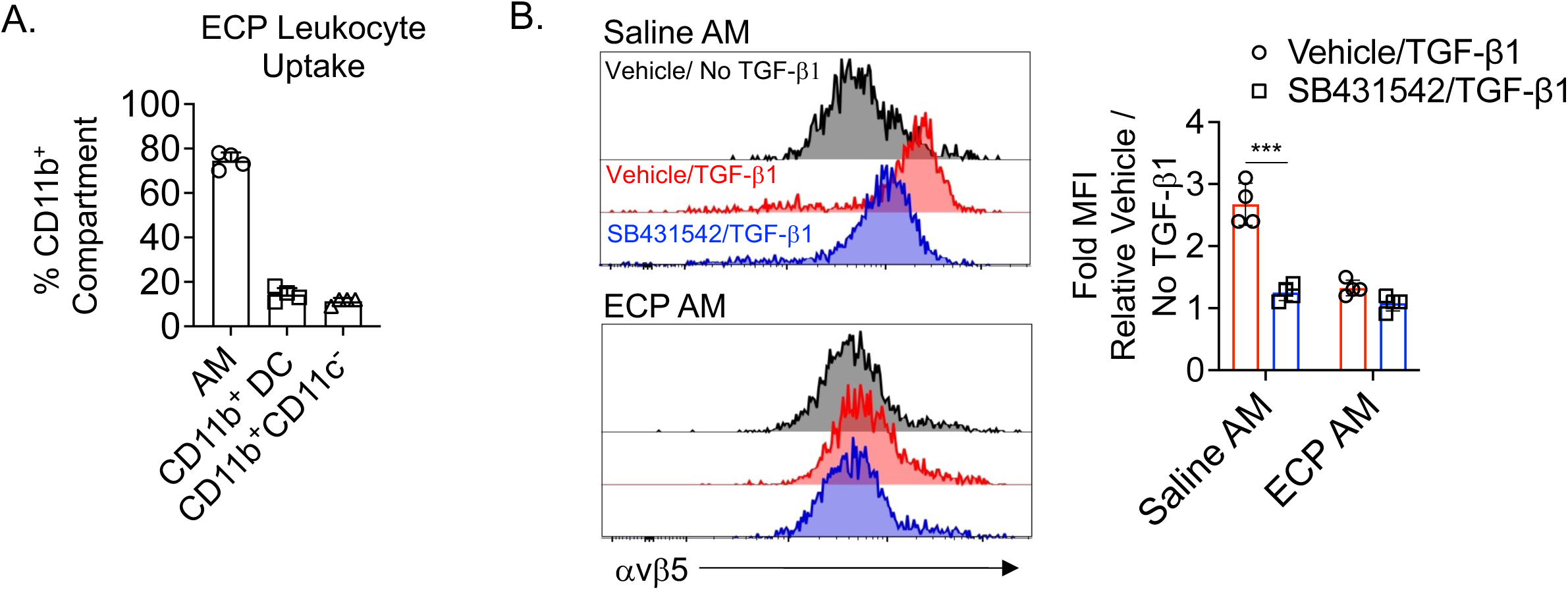
ECP-treated leukocytes are engulfed by AM, which in turn induces resistance to TGF-β mediated avβ5 upregulation. (A) CellTrace^633^-labeled ECP-treated leukocytes were injected into 3T-FVB lung recipients and analyzed for engulfment by the intragraft CD11b^+^ cell compartment. Data shown is the mean percent abundance ± S.D. for indicated phenotypes. (B) Saline and ECP-treated AM were pre-treated with 10 mM SB43152 or vehicle (DMSO) and then stimulated with or without TGF-β1. Data shown are representative FACS plots (N=4) and fold MFI changes in avβ5 expression normalized to respective non-TGF-β-treated DMSO-pretreated controls.

**Fig. S3:**
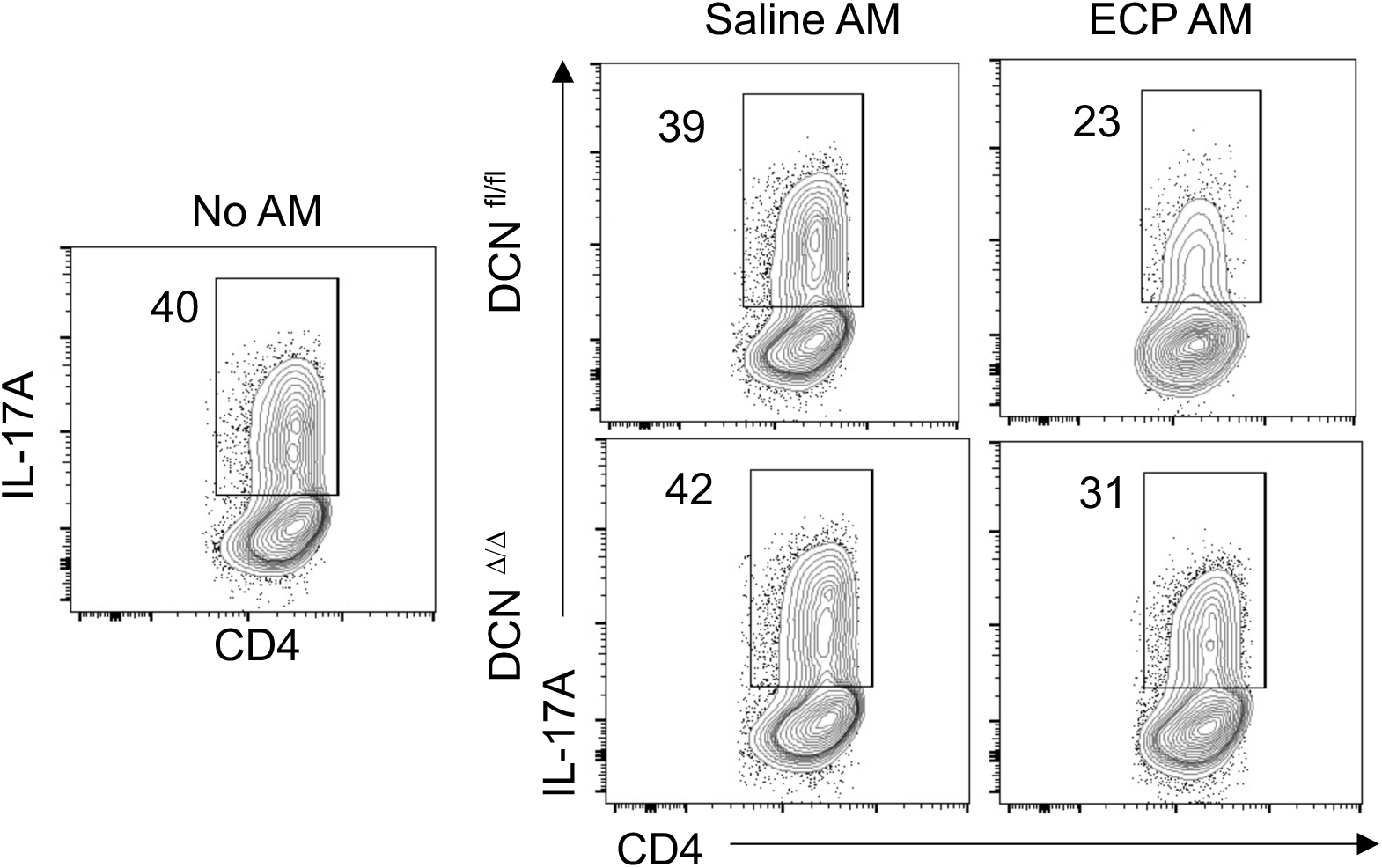
ECP-treated AM inhibits T_h_17 development in DCN-dependent manner. Representative FACS bivariate plots (N=5) of plate bound CD3ε and CD28 Ab-mediated stimulated B6 naïve CD4^+^ T cell cultured for 4 days in the presence or absence of indicated AM conditioned supernatants added in a 1:1 ratio to T_h_17 polarization medium that contains 10 ng/ml TGF-β1 and assessed for intracellular IL-17A expression.

**Fig. S4:**
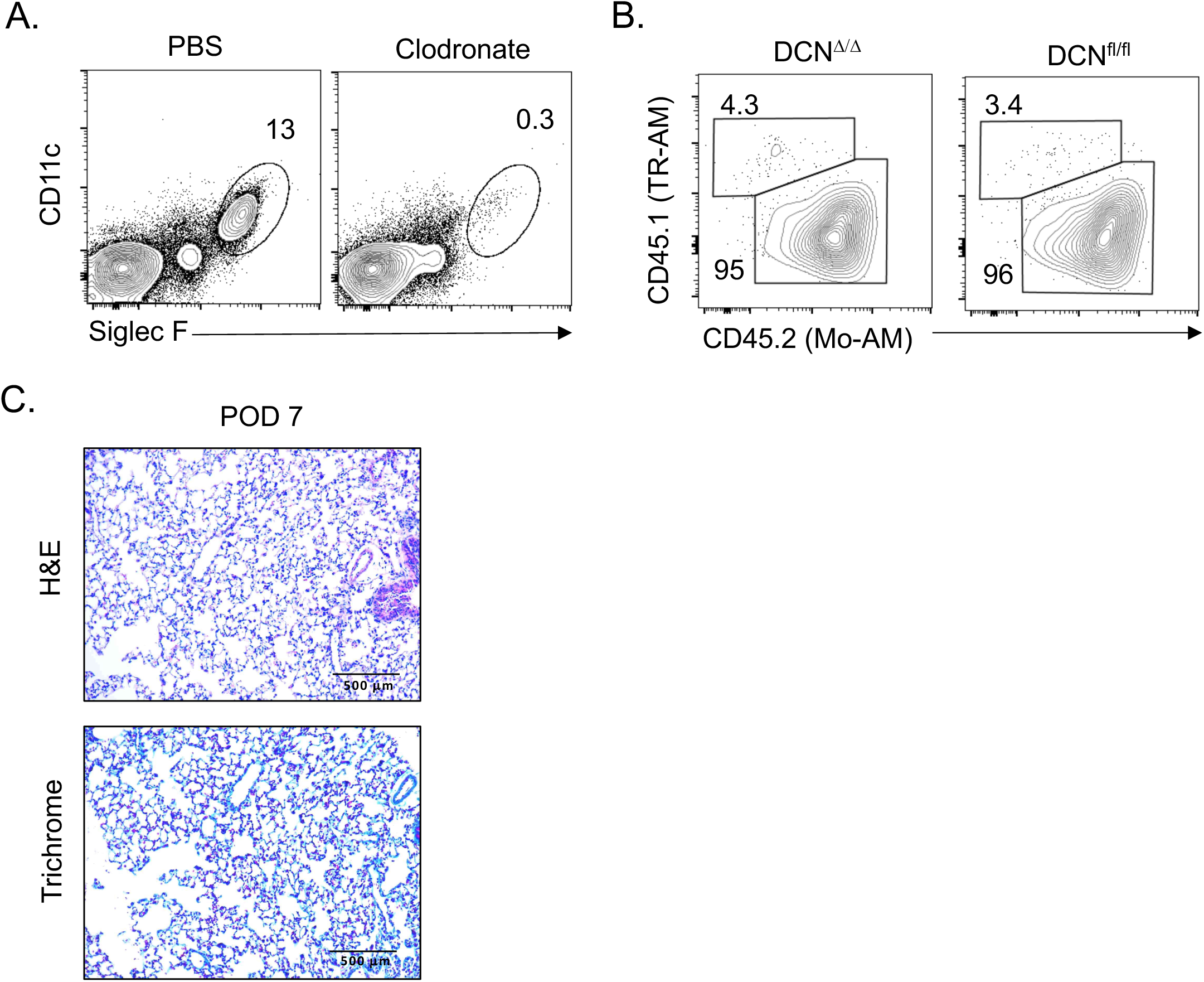
Donor lung clodronate TR-AM depletion and reconstitution with donor-derived AM does not spontaneously induce OB lesions. **(A)** A representative FACS plot of FVB lung donors 1 day after receiving 100 ul saline or clodronate liposomes. Data are representative of 2 independent experiments. Representative **(B)** lung allograft FACS plots and **(C)** histology from POD 7 DCN^Δ^*^/^*^Δ^ and DCN^fl/fl^ lung recipients of 3T-FVB allografts. Data are representative of at least 2 independent experiments.

**Fig. S5.**
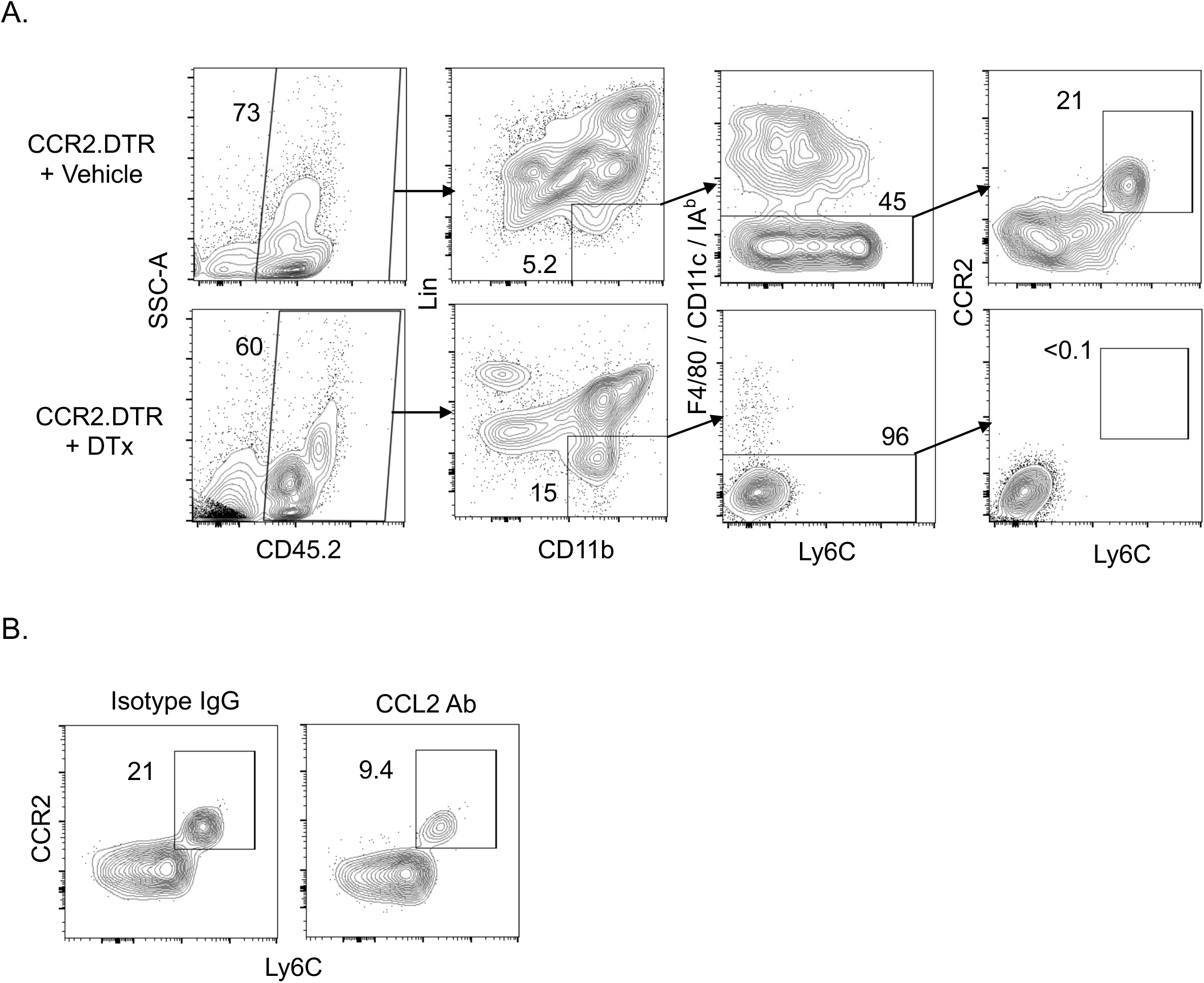
Depletion of monocytes from lung recipients. **(A)** FACS plot schema to measure allograft CCR2^+^ monocyte abundance of CCR2^DTR^ recipients 3 days after receiving PBS or 10 ng/g body weight of i.v. diptheria toxin. Data are representative of at least 2 independent experiments where Lin represents a cocktail of CD90.2, B220, NK1.1 and Ly6G-specific Abs. **(B)** 3T FVB allografts received 200 ug i.v. of CCL2 neutralizing Abs on POD 6, 9 and 12 and assessed for blood peripheral CCR2^+^ accumulation on POD 16. Data shown is representative from 2 experiments.

**Fig. S6.**
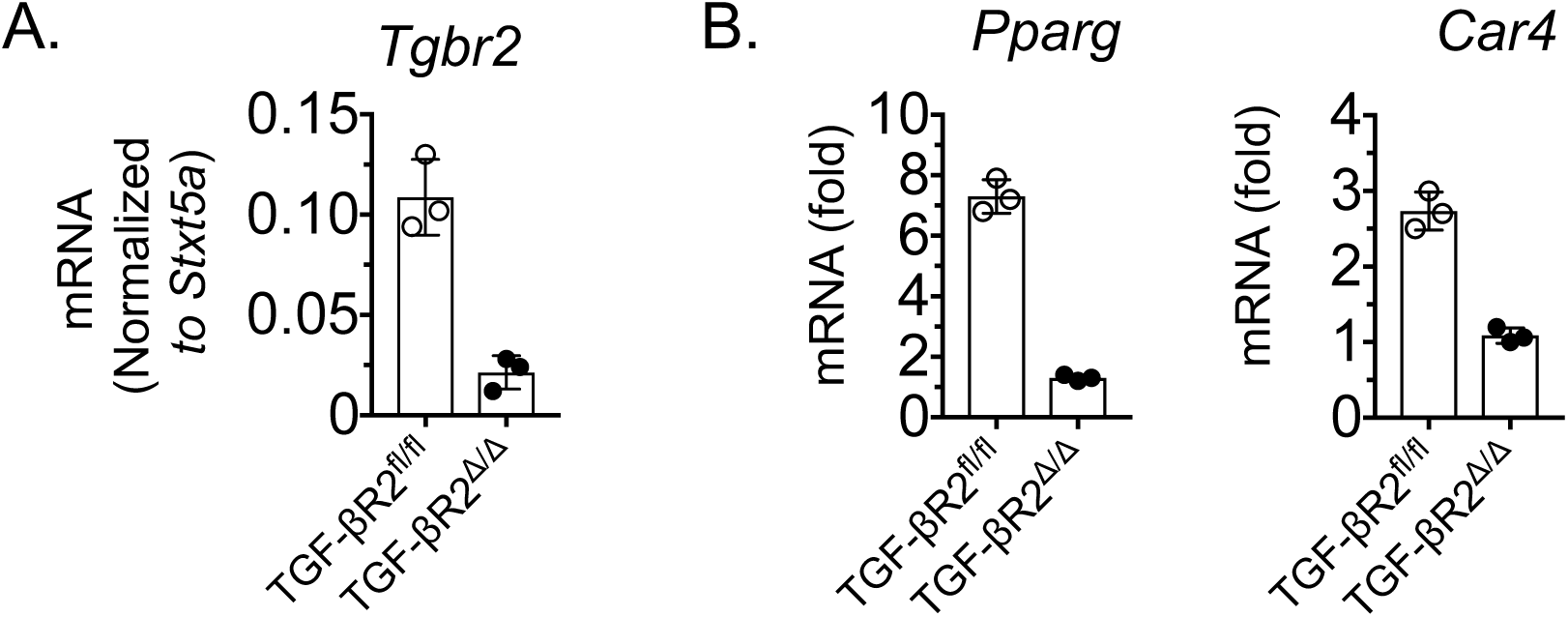
Validation of TGF-βR2 deletion in CCR2^+^ monocytes. (A) TGF-βR2^fl/fl^ and TGF-βR2^Δ/Δ^ mice were treated tamoxifen i.p. every other day for 10 days, rested for 5 days, FACS sorted for peripheral CCR2^+^ blood monocytes and assessed for (**A**) Tgbr2 transcript levels or (**B**) stimulated overnight with 5 ng/ml of TGF-β1 and assessed for *Pparg* and *Car4*. For (A) data shown is normalized to the macrophage housekeeping gene *Stxt5a* and (**B**) shown normalized to TGF-βR2^Δ/Δ^ levels where bars represent means ± S.D.

**Fig. S7.**
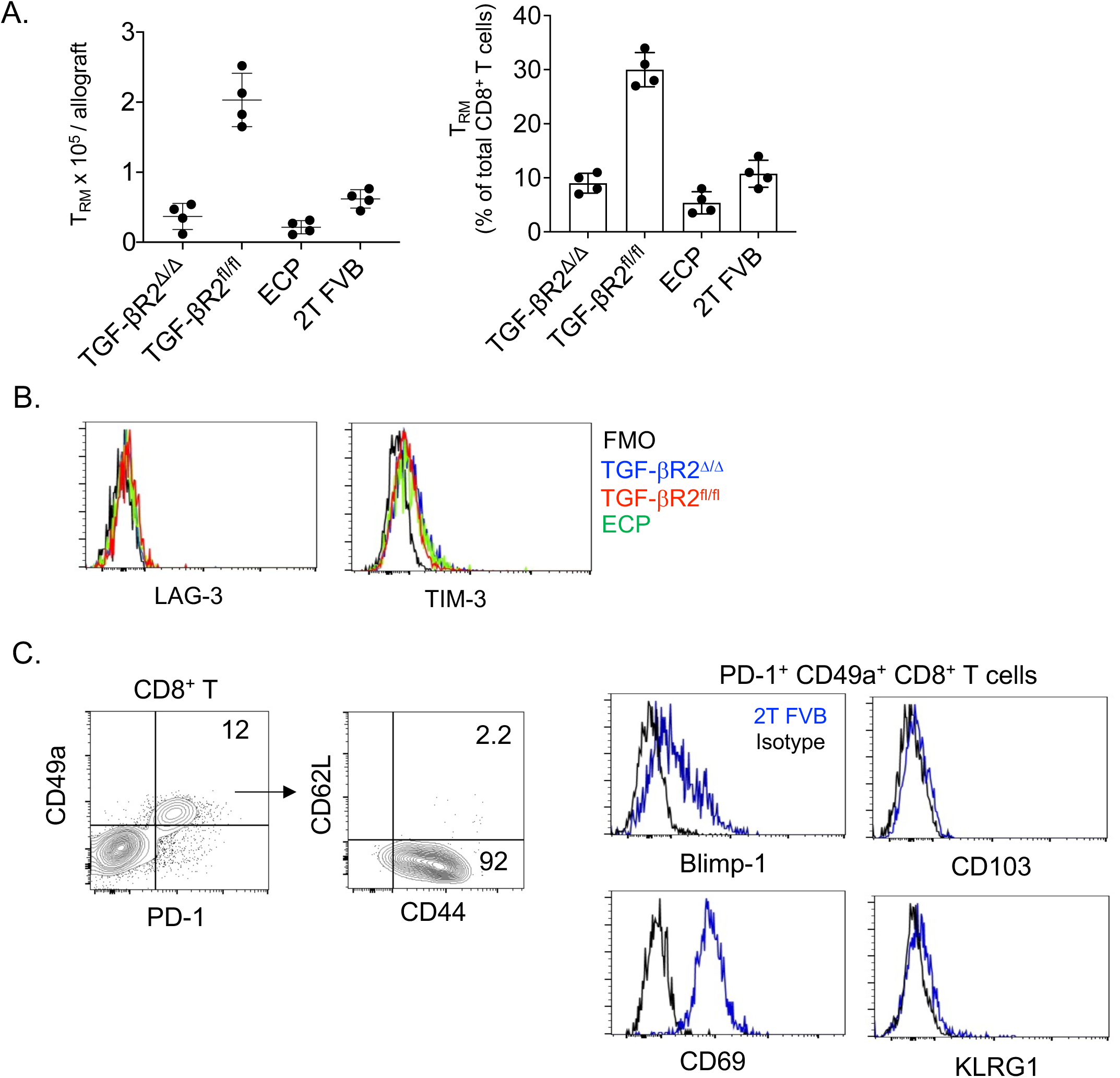
T_RM_ phenotype and abundance in lung transplants. **(A)** Intragraft percent abundance relative the whole CD8^+^ T compartment and intragraft numbers of T_RM_ for indicated transplant and treatment conditions. Bars represent means ± S.D. (**B**) Representative (N=4/group) FACS histograms of T_RM_ LAG-3 and TIM-3 expression. (**C**) Representative N=4/group) FACS plots and histograms of T_RM_ marker expression patterns found within 2T-FVB allografts.

**Fig. S8.**
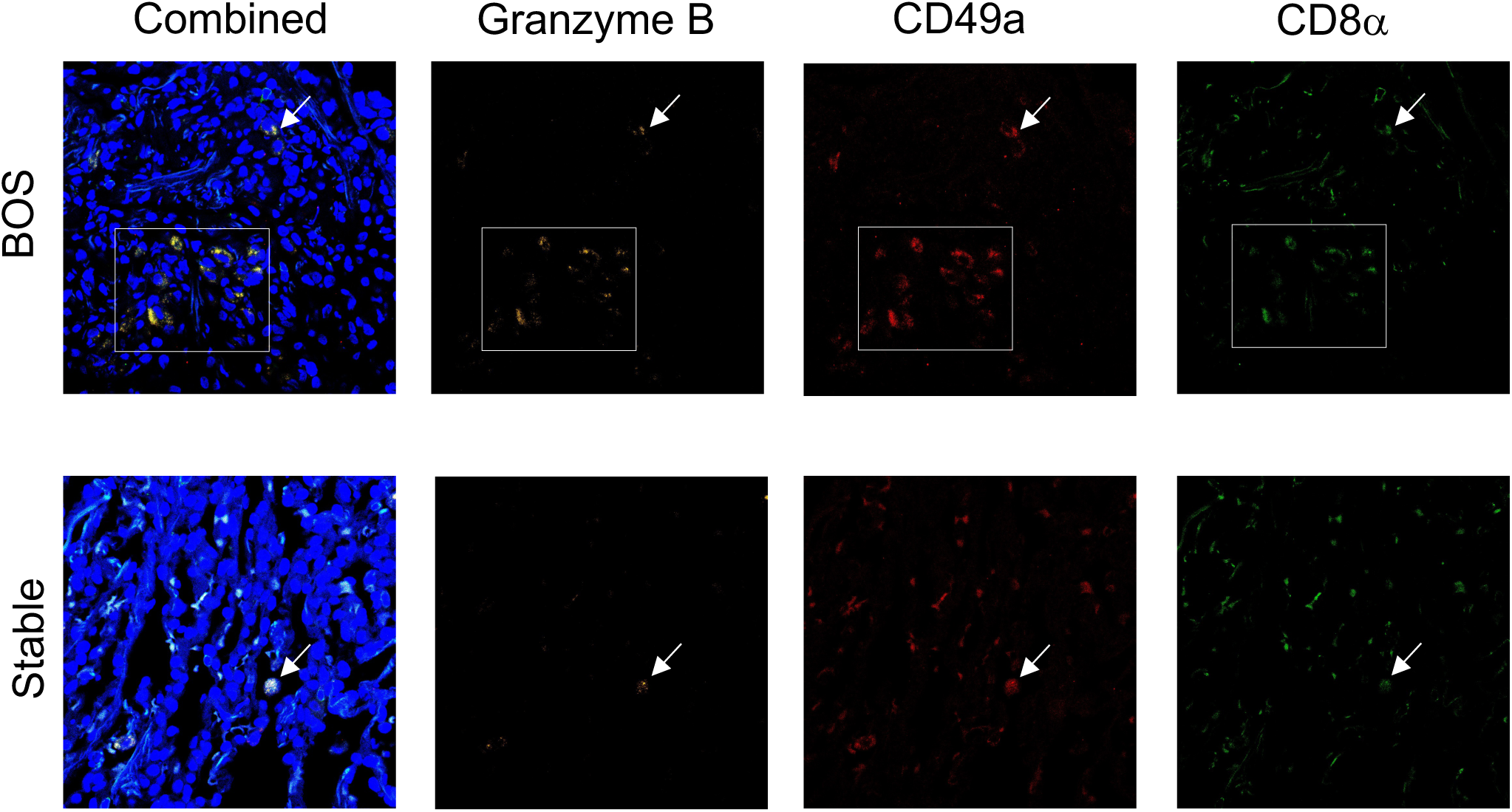
Clusters of Gzmb^+^CD49a^+^CD8^+^ T cells in BOS subjects. BOS explant tissue (BOS) and core biopsies from patients without evidence of CLAD or antibody-mediated rejection (Stable) were stained with indicated antibodies. Data are representative stains from BOS (N=4) and stable recipients (N=6). The white box represents a cell cluster and arrows show a single cell.

**Fig. S9.**
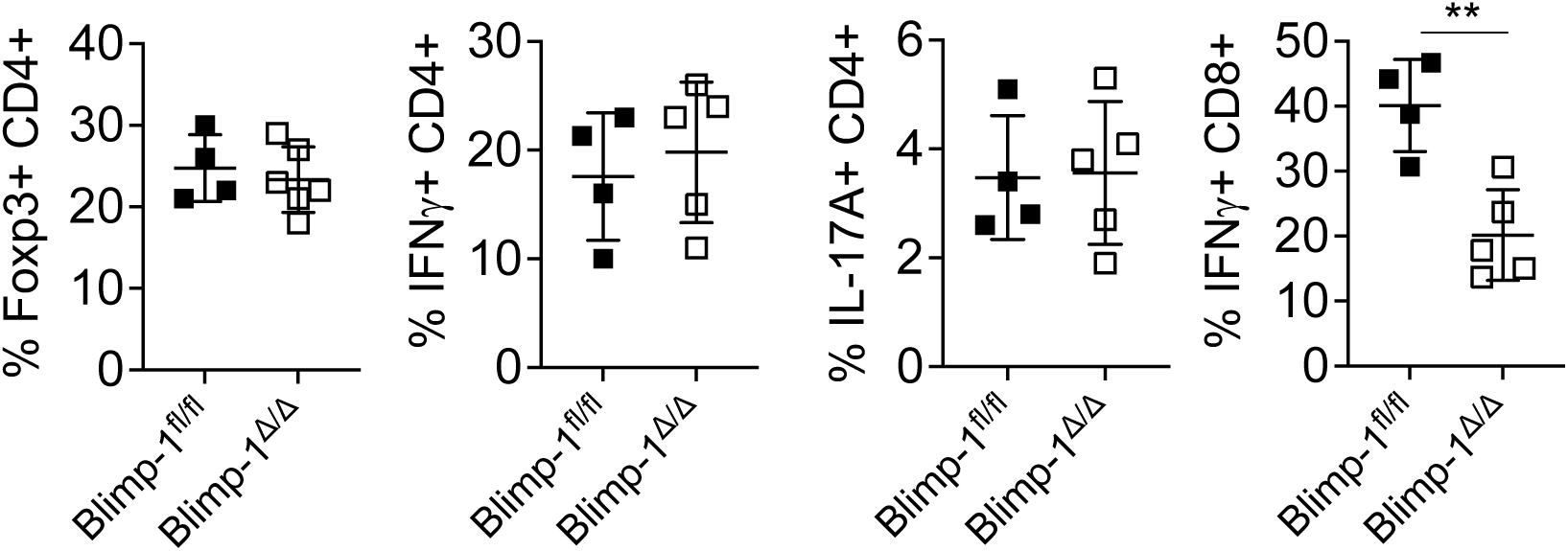
CD8^+^ T cell deficiency of Blimp-1 prevents reduces intragraft IFN-γ^+^ CD8^+^ T cell accumulation. Flow cytometric analysis of indicated intragraft effector T cells in Blimp-1^flfl^ and Blimp-1^Δ^*^/^*^Δ^ recipients. Bars represent means ± S.D where **p < 0.01.

